# Understanding systems level metabolic adaptation resulting from osmotic stress

**DOI:** 10.1101/2024.03.19.585265

**Authors:** Alexandre Tremblay, Pavlos Stephanos Bekiaris, Steffen Klamt, Radhakrishnan Mahadevan

## Abstract

An organism’s survival hinges on maintaining the right thermodynamic conditions. Osmotic constraints limit the concentration range of metabolites, affecting essential cellular pathways. Despite extensive research on osmotic stress and growth, understanding remains limited, especially in hypo-osmotic environments. To delve into this, we developed a novel modeling approach that considers metabolic fluxes and metabolite concentrations along with thermodynamics. Our analysis of *E. coli* adaptation reveals insights into growth rates, metabolic pathways, and thermodynamic bottlenecks during transitions between hypo- and hyper-osmotic conditions. Both experimental and computational findings show that cells prioritize pathways that have higher thermodynamic driving force, like the pentose phosphate or the Entner–Doudoroff pathway, under low osmolarity. This work offers a systematic and mechanistic explanation for reduced growth rates in hypo- and hyper-osmotic conditions. The developed framework is the first of its kind to incorporate genome wide constraints that consider both natural logarithm and actual metabolite concentrations.

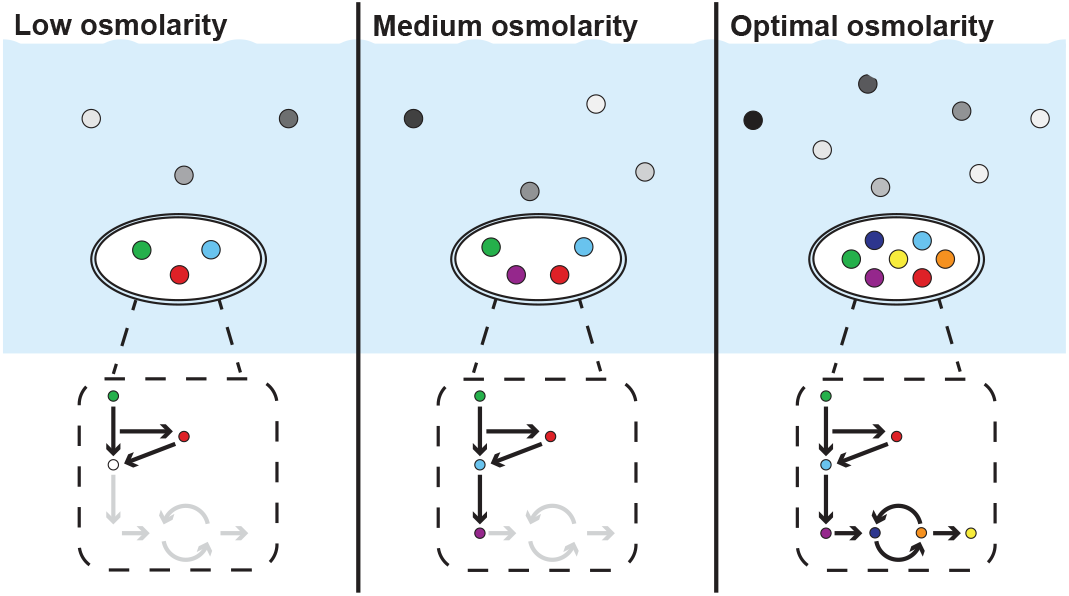

## Introduction

Microorganisms are found in nearly every corner of the earth, from hostile to fertile. Although most species have evolved to occupy a limited niche, some can maintain growth in diverse environments. *E. coli*, for example, a bacterium found in the guts of warm-blooded animals, can grow in anaerobic or aerobic conditions, at low and high temperatures, and in environments with high and low osmotic pressure. This remarkable adaptability implies that these generalists can maintain the internal conditions that keep biochemical pathways involved in maintenance and growth, feasible. Translated in chemical terms, feasibility means that these pathways remain thermodynamically driven, i.e., that the Gibbs free energy of each essential reaction remains negative in the direction of the required metabolites synthesis. However, this can be a difficult task under osmotic stress, where the internal metabolite concentrations have to be balanced with the cell’s environment to prevent plasmolysis [1].

In prokaryotes, changes in osmotic pressure cause water to flow passively through aquaporins (e.g., AqpZ) [2]. The passive nature of these transporters implies that hyper- or hypo-tonic environments can compromise the integrity of the cell wall and in extreme conditions could even lead to complete lysis. Cells can rapidly react to this stress by importing and producing osmolytes such as glutathione, potassium or trehalose [3–5] and by rapidly exporting metabolites through non-specific mechanosensitive channels such as MscL and MscS [6, 7].

Even after adaptation, the growth rate was found to remain impaired by the media’s osmolarity [8, 9]. This led to the hypothesis that turgor pressure might be required to trigger peptidoglycan biosynthesis and promote cellular growth. However, Rojas et al., 2014 showed that cell-wall elongation happens even during plasmolysis and thus that mechanical stress is not required for growth [10]. These results hint at some other mechanism at play that have yet to be identified. This gap in knowledge could be explained by the lack of a mechanistic understanding of the impact of osmolarity on the metabolism. This gap is particularly steep for hypo-osmotic environments, as most efforts has been directed toward hyper-osmotic stress.

In recent decades, constraint-based modelling with its core method flux balance analysis (FBA) has become a powerful tool for the analysis of the metabolism of microorganisms [11–13]. Extensions to the framework include the addition of thermodynamic constraints allowing, for example, the calculation of feasible metabolite concentration ranges [14–17]. Recently, ionic strength and osmotic balance alongside multiple other abiotic constraints associated with the extracellular environment were used to analyze the space of concentrations assuming fixed fluxes [18]. However, the framework developed still cannot be used to evaluate how the growth rate and metabolic fluxes are shaped by osmolarity.

One major issue with the integration of osmolarity with metabolic fluxes is that flux feasibility depends on the natural logarithm of metabolite’s concentration whereas osmolarity affects the total sum of metabolite concentration. Thus combining both makes the problem non-linear.

In this article, we introduce a novel formulation called *osmotically constrained Flux Balance Analysis* (ocFBA) which integrates metabolite concentrations, stoichiometry of reaction, osmolarity and thermodynamic feasibility to predict the growth phenotype of *E. coli* throughout osmotic stress. To get around the problem’s non-linearity, a series of linear constraints were introduced to approximate metabolite concentrations from their natural logarithm with high accuracy. Using ocFBA, we were able to calculate how the growth phenotype of *E. coli* is shaped by osmotic limitations, and identified thermodynamic bottlenecks reactions behind these limitations. These findings were subsequently validated through growth assays and metabolome analysis. This study constitutes, to the best of our knowledge, the first analysis relating osmotic pressure to growth rate in genome-scale models and the first to provide an experimentally validated theoretical framework explaining growth impairments measured at low osmolarities.

## Results

### The ocFBA framework

To evaluate the impact of osmolarity on the metabolism, we developed a framework, ocFBA, that allows for the calculation of metabolic fluxes in environments with different osmotic pressure.

There are four interconnected levels of constraints in ocFBA: a first one containing the classical FBA constraints, a second one that ensures all reactions with non-zero fluxes are thermodynamically feasible, a third one that approximates the concentration of metabolites involved in active reactions and a fourth one that enforces osmotic constraints by limiting the total maximum concentration of metabolites (see Methods section for full details). The first two levels of constraints are useful to calculate the thermodynamic driving force of metabolic reactions as outlined in the studies describing Max-min Driving Force (MDF) [19] and its extension, OptMDFpathway [20]. Taken together, these constraints ensure that the calculated optimal solution satisfies osmotic constraints and is thermodynamically feasible, i.e. the metabolic pathways maintain a threshold level of driving force (MDF), while maximizing the growth rate. The imposition of osmotic and thermodynamics constraints along the ability to maximize growth is original to this work.

Typically, thermodynamic feasibility is calculated from the natural logarithm of the concentration of metabolites. However, according to the van’t Hoff formula (equation 8), osmotic pressure is a function of the water activity, which is itself a function of the total molar concentration of components in solution (equation 14). Thus, the third level of constraints approximates the concentration of metabolites from their natural logarithm through a piecewise linear function (equation 21). The fourth level acts directly on these estimated concentrations, forcing the sum of all metabolites involved in the solution to remain lower than a defined threshold, represented by the variables Φ and *θ*. Φ factors in the effect of ions, both in the intracellular and extracellular environment as well as the total concentration of metabolites in the extracellular environment. As such, it can be seen as a proxy for the osmolarity of the extracellular environment. *θ* represents the maximum tolerable osmotic pressure exerted on the cell wall. Typically, *E. coli* maintains a negative osmotic pressure of 0.3 atm [21], although negative pressure is not required for growth [10]. In our formulation we allow for the pressure in the cell to go from -0.3 to 0 atm using Φ and *θ* (equation 19). The units of Φ and *θ* are mol/L. Overall, the resulting constraint added to the optimization problem is represented by equation 20 (see Methods).

In addition to the constraints, to simulate the increased energy maintenance demand at higher osmolarity caused by potassium importation and osmoprotectant production [3–5], a hyper osmotic condition constraint was added. This constraint includes a pseudo-reaction called Osmotic maintenance Demand (OmD), which leads to the consumption of ATP, proportionally to Φ, upon reaching a Φ_*threshold*_ value.

Taken together, these modules constitute the basis of the optimization problem behind ocFBA, which is represented in equation (26). The framework of ocFBA can be used to calculate the maximal growth rate (*μ*) of a strain in an osmotically constrained environment.

### Toy model representation

To illustrate how the solution changes with Φ, a proxy for the osmolarity of the extracellular environment, the optimization problem was tested on a toy model made up of 4 reactions (Fig. 1). On this smaller network, it can be seen that lower Φ values reduce the pool of feasible reactions, forcing the problem to generate biomass preferably through smaller networks, despite their lower biomass yield. As Φ increases, more reactions become feasible, since the minimal concentration of metabolites required to make these reactions thermodynamically feasible is met. In turn, the solution space increases, allowing for new pathways with higher ATP yields and therefore with higher growth rates.

**Fig 1.**
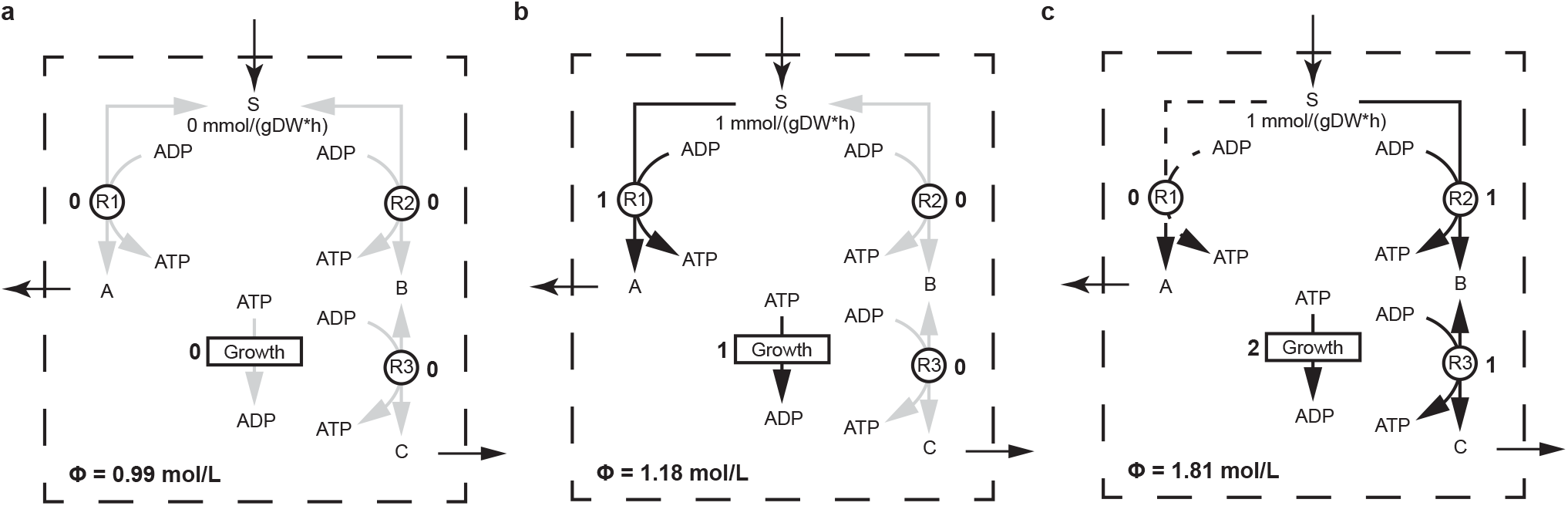
Example of how the osmotic concentration coefficient (Φ) affects the feasibility of reactions and max growth rate on a toy model. Feasible and infeasible reactions are shown by black and gray arrows, respectively. The number next to the reaction name indicates the flux computed by the algorithm. Dashed arrows represents feasible reactions that are not part of the optimal solution. All reactions are unbounded except for substrate (S) uptake which had a lower bound of -1. Reversibility is indicated by the direction of the arrows. For all reactions, the minimal and maximal metabolite concentration was set to 0.1 and 1 mol/L, respectively. Furthermore, the standard Gibbs free energy Δ_*r*_*G*^*′°*^ was set 3 kJ/mol. **a)** ocFBA with Φ = 1.00 mol/L. **b)** ocFBA with Φ = 1.19 mol/L. **c)** ocFBA with Φ = 1.82 mol/L

### Growth under osmotic stress

Maintaining growth in environments with low osmolarity can be particularly difficult, as cells are forced to suitably utilize their resources to prevent water from flowing in and out of the cell. To evaluate the impact of osmolarity on the growth *E. coli*, several ocFBAs (equation 26) were performed with increasing osmotic concentration coefficient (Φ) values (Fig. 2a) using the genome-scale model *i* ML1515 [22]. To ensure that every solution were thermodynamically driven and hence feasible, the minimal driving force across all active reactions value (B) was set to 0.01 kJ/mol.

**Fig 2.**
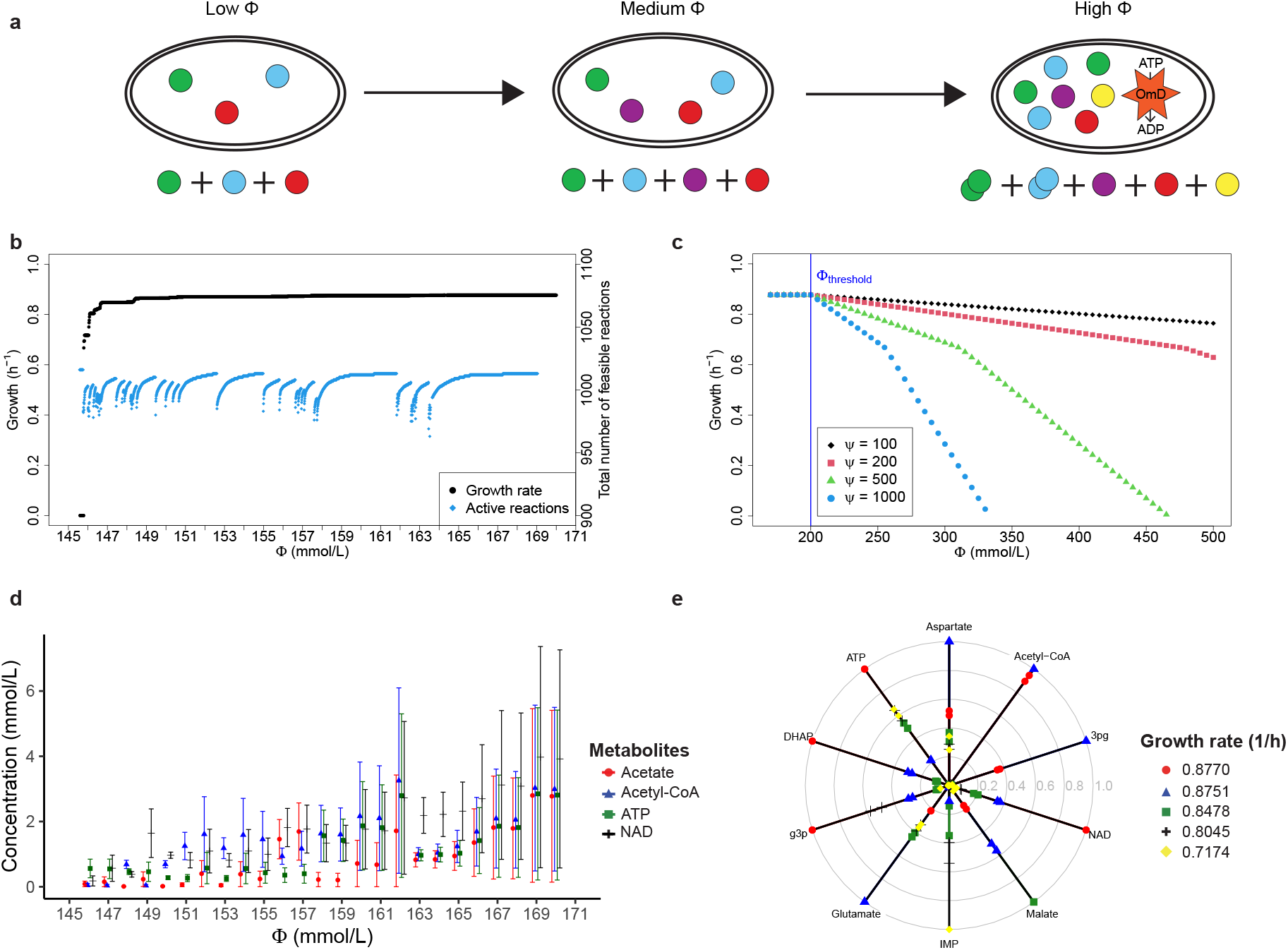
Effect of the osmotic concentration coefficient (Φ) on maximal growth rate and metabolite concentration. All results were calculated using the genome scale model of *E. coli i* ML1515. **a)** Schematic of the analyses performed. Φ affects the maximum sum of all metabolites. Increasing Φ allows for a higher abundance of metabolite, which in turn affects the maximal number of feasible reactions in the solution **b)** Impact of Φ on the maximal growth rate (black) and on the total number of thermodynamically feasible reactions at that maximal growth rate (cyan). The total number of reactions involved in the thermodynamic constraints is 2266, but the maximal number of simultaneously feasible reactions is 1017. **c)** Impact of the hyper-osmotic constraint on growth for 4 different *ψ* values (100, 200, 500 and 1000 mmol *·* L/(mol *·* g-DW *·* h)). The threshold (Φ_*threshold*_) at which the hyper-osmotic constraint is activated was set to 0.2 mol/L. **d)** Feasible concentration range of acetate (red circle), acetyl-CoA (blue triangle), ATP (green square) and NAD (black cross) at the maximal growth rate for different values of K. **e)** Concentration range for the minimal value of *ϕ* needed to reach a growth rate of 0.8770*h*^−1^ (red circle), 0.8751*h*^−1^ (blue triangle), 0.8478*h*^−1^ (green square), 0.8045*h*^−1^ (black cross), 0.7174*h*^−1^ (yellow diamond). Metabolites are presented on a proportional scale, where the highest calculated concentration is 1 and the smallest is zero. The center of the figure represents the zero point. The minimal and maximal values are represented by two points of the same color and shape.

This method was used to evaluate the effect of changing Φ on the growth rate *μ*. The results show that the growth rate of *E. coli* increases in stepwise fashion as Φ increases (Fig. 2b). Growth is initially infeasible until Φ reaches about 0.1458 mol/L. From that point on, it sharply increases before plateauing briefly around 0.7174 h^-1^. All subsequent growth improvements after this initial increase are progressively smaller. The maximal growth rate plateaus several more times after this point before reaching 0.8770 h^-1^, the maximal growth rate calculated using regular FBA without any osmotic constraints.

For each value of Φ, the maximal number of reaction with non-zero flux with an assigned Gibbs free energy values was calculated by fixing the value for *μ* and maximizing the number of feasible reactions instead of *μ* in equation (26). It was found that the maximal number of feasible reactions appears to be tightly related to the growth rate plateaus (Fig. 2b). For each growth plateau, increasing Φ increases the number of feasible reaction. However, as Φ increases, the maximal feasible *μ* increases and the maximal number of feasible reaction at that growth rate may diminish. For example, from Φ = 0.1467 mol/L until Φ = 0.1475 mol/L, growth remains constant at 0.8478 h^-1^. However, the number of feasible reactions increases from 985 to 1009. As soon as the maximum growth increases to 0.8479 h^-1^, the number of feasible reactions goes down to 987 for this particular growth rate. This results points towards specific bottleneck reactions which are required for greater growth values. The maximal number of simultaneously feasible reactions (1017) is achieved when Φ reaches 0.55 mol/L. However, no further growth improvements are measured past 0.1643 mol/L.

The impact of the osmotic maintenance demand reaction (OmD) on growth was evaluated by setting Φ_*threshold*_ to 0.2 mol/L and *ψ* to different values (Fig. 2c). This threshold was chosen somewhat arbitrarily, as it simply needed to be higher than the value after which Φ no longer constrains the growth rate. Φ_*threshold*_ and *ψ* should be chosen to fit experimental results. The results shows that the effect of OmD on growth can be divided into two separate linear fragments. The first linear fragment is between 0.8770 h^-1^ and 0.6684 h^-1^, and the second one is between 0.6684 h^-1^ and 0 h^-1^. Increasing *ψ* increases the sharpness of the slopes.

The feasible metabolite concentration range associated with each growth phenotype was calculated for different values of Φ through a concentration variability analysis (CVA) (Fig. 2d) [23]. Within the ocFBA framework, CVA calculates the maximum and the minimum concentration a metabolite can have for a specified growth rate, minimal driving force (*ϵ*) and Φ value. To evaluate the general change in concentration of metabolites in different osmotic conditions, the analysis was narrowed to four metabolites: acetate, acetyl-CoA, ATP and NAD. The former two were chosen because their abundance is known to fluctuate at different stages of growth [24, 25] and the latter two were chosen because they are central co factors.

The results show that, between 0.146 and 0.147 mol/L, the feasible range of acetyl-CoA and acetate concentration remains extremely limited, tightly constrained around the lower bound of the metabolite concentration. From 0.147 mol/L onward, acetyl-CoA’s concentration range remains well above the minimal concentration, indicating that it is now required in greater concentrations. For acetate, the situation is similar. However, its abundance appears to be tied to specific growth rates rather than to universal reactions required to achieve a growth rate past a certain threshold. This can be seen for the range of 0.156 until 0.157 mol/L and from 0.163 mol/L onward, where acetate’s lower concentration remains above the minimum. As for ATP and NAD, their concentrations remained well above the minimal allowable concentration throughout the studied range.

In general, the range of metabolite concentrations appears to be more constrained at the beginning and between plateaus. This result once again points toward specific bottlenecks that remain infeasible until Φ is high enough to allow for metabolite concentrations that meet their thermodynamic demands. Until this happens, the range of concentration for every metabolite will continue to increase, as long as it does not reduce the value of the optimal solution. Using the toy model for comparison (Fig. 1), plateaus represent states of the metabolome between Φ = 1.18 and Φ = 1.81 mol/L. Until metabolites can accumulate to levels that make R2 and R3 feasible, the growth rate remains constant, but the feasible concentration of all metabolites can increase.

Thus far, growth appears to be tied to reactions whose feasibility depends on the accumulation of specific metabolites. To identify these metabolites as well as understand how their abundance changes with the rate of biomass formation, the concentration range of every metabolite was calculated for five osmotic concentration coefficients: Φ = 0.1458, 0.1461, 0.1467, 0.1512, and 0.1643 mol/L (full list of metabolite and their concentration range available on GitHub). These represent the lowest values of Φ at the beginning of the plateaus associated to the growth rates 0.7174, 0.8045, 0.8478, 0.8751 and 0.8770 h^-1^, respectively.

The majority of the metabolites studied (717/783 at 0.8770 h^-1^) had 1 *μ*M as their maximal feasible concentration. Coinciding with the minimal concentration allowed in ocFBA, these metabolites either need to be maintained in low concentrations to make a reaction feasible or are not involved in the thermodynamic feasibility sub-problem.

A total of 76 metabolites were found to have a minimal concentration above the lower bound. Only the relative abundance of a selected few, whose behaviours were found to be representative of the different trends observed, were plotted (Fig. 2e). We found that generally (42/71), the basal metabolite concentration tends to be higher at 0.8770 h^-1^. Some metabolite’s basal concentration remained unchanged throughout the analysis. These include citrate and D-glucosamine 6-phosphate. These metabolites are likely associated with a single group of essential reactions. The rest of the metabolites, such as malate, appeared tied to a specific phenotype.

### Bottlenecks and osmotic balance

To further understand how and what reactions limit the solution space at lower Φ values, thermodynamic bottleneck reactions were calculated by finding all reactions whose driving forces were equal to *B* in equation (26) (see methods section on thermodynamic bottlenecks calculation) (Table 1). The same *μ* and Φ values as in the previous analysis were used.

**Table 1.**
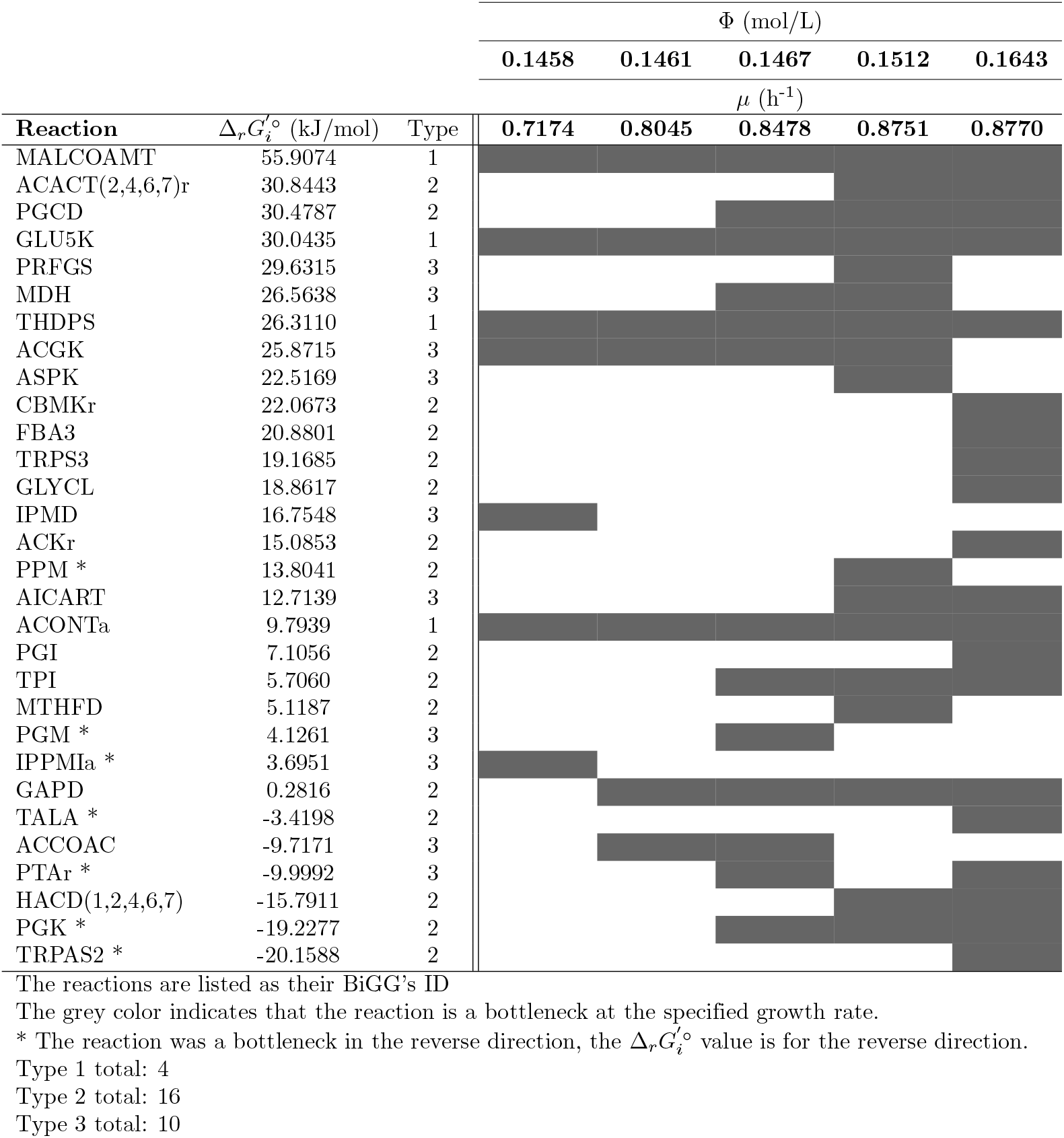
Thermodynamic bottlenecks (kJ/mol) with the lowest values of Φ (mol/L) allowing specific growth rates.

Our osmotic constraint forces cells to utilize metabolites parsimoniously, leading to distinct sets of bottlenecks. However, some essential reactions (growth cannot occur without them) with high 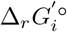 remain constraining regardless of Φ. In our table, these were defined as type 1 bottlenecks. This was the case for malonyl-CoA methyltransferase (MALCOAMT), glutamate 5-kinase (GLU5K), tetrahydrodipicolinate succinylase (THDPS) and aconitase (ACONTa). Metabolites associated with these reactions are likely to be more abundant. It is notably the case of glutamate (involved in GLU5K), whose concentration is consistently above the minimal allowable concentration in Fig. 2d.

The rest of the osmolarity-based bottlenecks can be subdivided into two categories: reactions that are required to grow at high rates (type 2) and reactions that are always part of the solution but are only bottlenecks in the presence of other competing reactions (type 3). Across all tested phenotype, a total of 30 reactions were identified as bottlenecks. Type 2 bottlenecks were the most common (15/30), followed by type 3 (10/30) and by type 1 (4/30).

In the type 2 category of bottlenecks, we find the acetyl-CoA C-acyltransferase reactions (ACACT7r, ACACT6r, ACACT4r and ACACT2r). They have the second highest standard Gibbs free energy out of all bottlenecks and are required to reach growth rates higher than 0.8698h^-1^. Since all of these reactions rely on acetyl-CoA as their substrate, this phenomenon also explains why higher concentrations of acetyl-CoA are found when Φ reaches 0.147 mol/L (Fig. 2d).

In the type 3 category of bottlenecks, we find the reaction acetyl-CoA carboxylase (ACCOAC). ACCOAC is always required in each of the studied phenotypes. Despite its negative Δ*G*^′°^value. ACCOAC is a bottleneck when Φ = 0.1461 and Φ = 0.1467 mol/L. This reaction leads to the conversion of acetyl-CoA and bicarbonate into malonyl-CoA through the dephosphorylation of ATP. Malonyl-CoA is the substrate of the essential reaction malonyl-CoA methyltransferase (MALCOAMT). Having the highest Δ*G*^*′°*^ value of all bottlenecks, MALCOAMT relies on high concentrations of malonyl-CoA to be feasible. In turn, this makes ACCOAC a bottleneck as well. ACCOAC is not a bottleneck at lower Φ values, notably due to the absence of glyceraldehyde-3-phosphate dehydrogenase (GAPD) from the list of active reaction at the maximal growth rate. Without GAPD, phosphate concentration can be maintained at lower concentrations. Since dephosphorylaton of ATP results in phosphate, lower concentration of phosphate makes ACCOAC more driven. For higher Φ values, the absence of ACCOAC from the bottleneck list can be explained by the greater flexibility associated to these conditions, allowing for higher bicarbonate concentrations.

About 1/5 of the bottlenecks (6 out of 30) had a negative 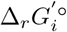. This is counter-intuitive as highly driven reactions with negative 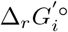 are typically expected to be feasible at lower substrate concentrations. However, given that multiple metabolites can be involved in many reactions, this kind of systematic network-based analysis instead of relying on 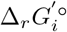 is the only way to correctly assess what reactions are bottlenecks and highlights the value of ocFBA.

### The reaction space under the osmotic constraint

To help visualize the solution space, Escher maps of the central carbon metabolism [26], representing the feasibility of each reaction for different values of Φ, were created (Fig. 3). The maps replicate what is demonstrated in Fig. 1, albeit at a much larger scale. Thus, the same underlying phenomenon explains both figures: Φ affects the pool of feasible reactions, which in turn affects the feasible max growth rate.

**Fig 3.**
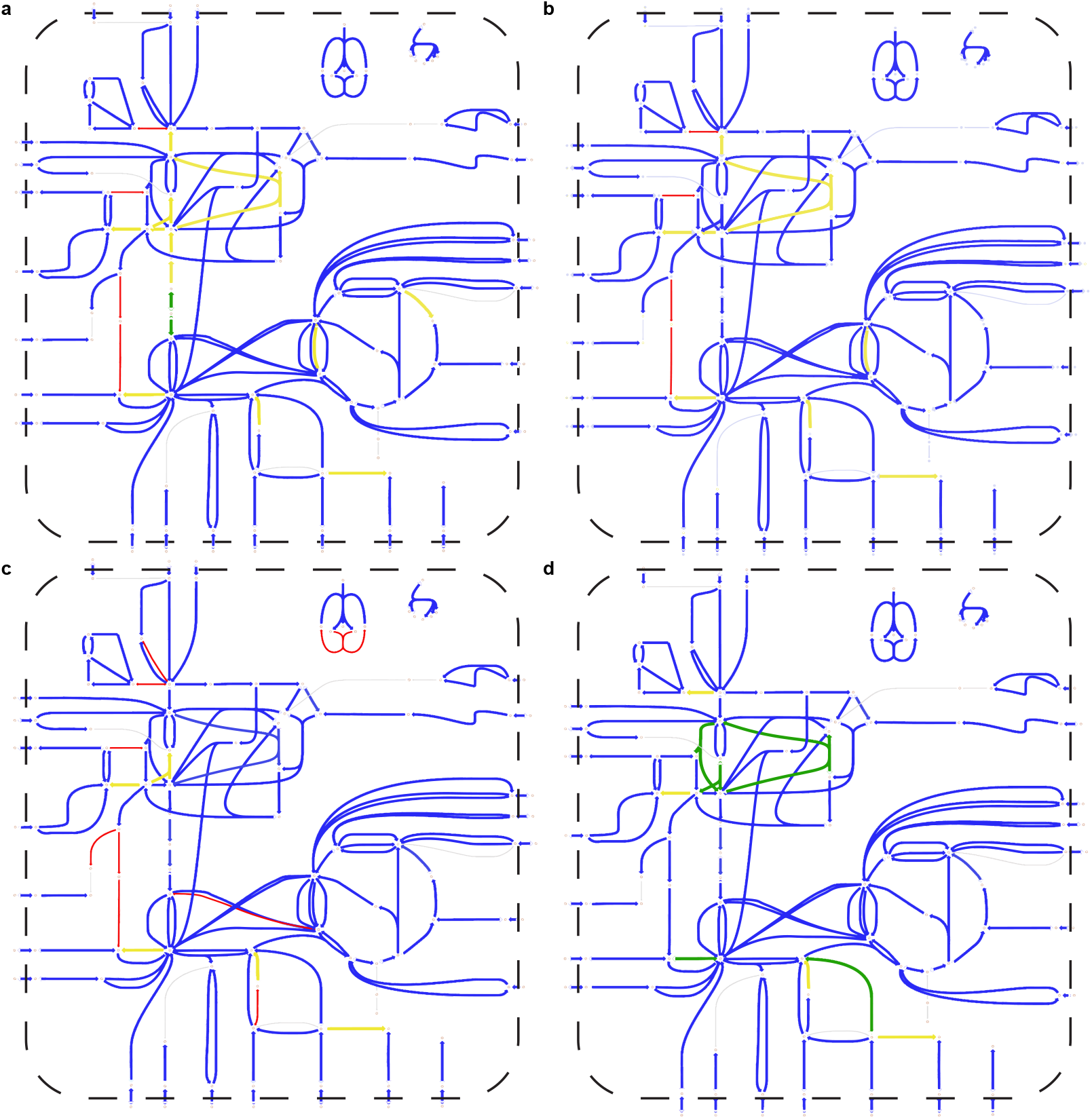
Feasibility of reactions in the central carbon metabolism of *E. coli* for different values of Φ. Each line represents the flux of each reaction and each dot represents a metabolite. Red lines mean that flux is never feasible, a blue line means that flux is only feasible in the forward direction, a yellow line means the reaction is only feasible in the reverse direction and a green line means that the reaction is feasible in both directions. Fluxes were calculated using the *E. coli* genome-scale metabolic model iML1515. **a)** Feasible fluxes when Φ = 0.1458 and *μ* = 0.7174^−1^. **b)** Feasible fluxes when Φ = 0.1461 and *μ* = 0.8045^−1^. **c)** Feasible fluxes when Φ = 0.1467 and *μ* = 0.8478^−1^. **d)** Feasible fluxes when at max growth rate (*μ* = 0.8770^−1^) and when osmolarity no longer limit growth (Φ = 0.1971).

Analysis of the feasible reactions in the central carbon metabolism of *E. coli* reveals that the osmotic constraint severely limits the pathways through which biomass can be produced. When Φ = 0.1458 mol/L and *μ* = 0.7174 h^-1^, glycolysis cannot proceed downstream towards pyruvate and the full citric acid cycle is not feasible. In turn, cofactor production and proton gradient generation must rely further on the Entner-Doudoroff (ED) and the pentose-phosphate (PP) pathways. These pathways have lower ATP yields but are a metabolite and enzyme efficient alternatives to glycolysis [27]

As the osmotic constraint becomes less limiting, more options become available. At Φ = 0.1461 mol/L and *μ* = 0.8045 h^-1^, glycolysis is almost feasible but it remains limited by glucose-6-phosphate isomerase (PGI), which is still only feasible in the reverse direction. However, the citric acid cycle is now feasible in the forward direction. At Φ = 0.1467 mol/L and *μ* = 0.8478 h^-1^, PGI becomes feasible in the forward direction but fructose-bisphosphate aldolase (FBA) is no longer feasible in the forward direction. Thus, to proceed towards pyruvate, both or either the ED and PP pathways need to generate glyceraldehyde 3-phosphate. The feasibility of reactions changing with Φ highlights that committing to a pathway in a resource-limited environment can mean eliminating other pathways due to thermodynamic bottlenecks.

As Φ increases and no longer limits growth, cells can achieve their maximal growth rate (*μ* = 0.8770 h^-1^). At this osmolarity, glycolysis, the citric acid cycle, the pentose phophate pathway and the Entner-Doudoroff pathway are all simultaneously feasible. This does not mean that the cell will utilize all of them, simply that it could allocate a fraction of its metabolome to these pathways.

### Experimental validation

Three main results from the computational analysis were subject to experimental validation: 1. the relation between the maximal growth rate and osmolarity, 2. the increased reliance on the ED and PP pathways at lower osmolarities and 3. the relation between the size of the metabolome and osmolarity. To do so, growth assays and metabolome analysis were performed using *E. coli* MG1655 or strains containing knock-outs preventing flux through the ED or PP pathway.

To validate the predicted growth kinetics presented in Fig. 2b) and c), a series of growth assays were done in our low osmolarity media supplemented with various concentrations of either NaCl or KCl (Fig. 4a). Two salts were used to ensure that the measured impact was not caused by the salt itself. We found the growth rate increases quadratically with the salt concentration in the media (R^2^ of 0.7457 for NaCl and R^2^ of 0.6105 for KCl). According to the results, the maximal growth rate initially increases with the salt’s concentration before plateauing with 0.1 mol/L of salt (theoretical osmolarity of 273 mOsm). After this point, the maximal growth rate starts decreasing. These results consolidate the literature on the topic, which shows similar kinetics and found that optimal osmolarity for growth to be around 300 mOsm [28–30].

**Fig 4.**
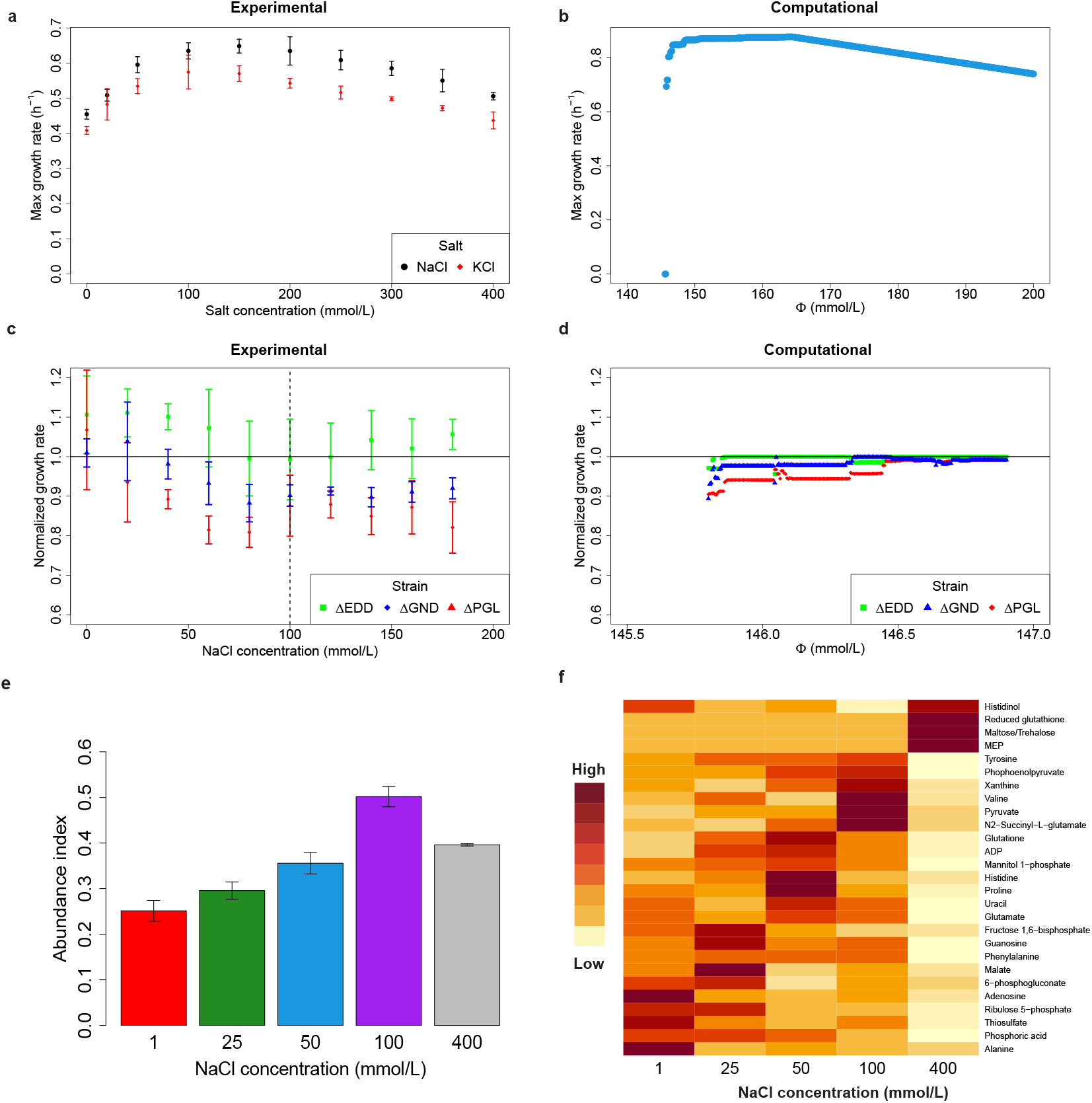
Growth and metabolome of different *E. coli* strains grown in media with various salt concentration. The theoretical osmolarity of the media without salt is 73 mOsm. **a)** Average maximal growth rate of *E. coli* MG1655 in our minimal media supplemented with NaCl (black dot) or KCl (red diamond). A total of 6 technical replicates were done for each condition (n=6). The error bars are the standard deviation associated with each treatment. **b)** Simulated growth rate when applying both the hyper- and hypo-osmotic constraints. **c)** Normalized maximal growth rate of JW1840 (ΔEDD, green), JW2011 (ΔGND, blue) and JW0750 (ΔPGL, red) subjected to different concentration of NaCl. To normalize, the average growth rate of the test strain was divided by the average growth rate of the control strain (JW1693) grown on the same plate. The error bars are the empirical uncertainty associated to each measurement calculated with the 3 technical replicates (see Methods section) (n=3). The dashed line represents the salt concentration leading to the highest growth rate, and the full line represents the normalized growth of the control. **d)** Predicted normalized growth for different Φ of three strains: ΔEDD (green) and ΔGND (blue) and ΔPGL (red) using ocFBA. The predicted growth rate of the strain was divided by the predicted growth rate of the wild-type. The black line represents the normalized growth rate of the wild-type strain. **e)** Metabolite abundance index for *E. coli* cultures subjected to different concentrations of NaCl using peak areas obtained with C18 column. The height of the bar represents the midpoint between the abundance index calculated using the lowest peak area measured with either technical replicate (n=2) and the highest peak area measured with either replicate. The error bar represents the distance from the midpoint to the minimal and maximal abundance index. Each measurements come from distinct samples. **f)** Heat map of the average metabolite concentration for each treatment for metabolite with high treatment variability. Darker shades mean that the average area of a metabolite associated with the treatment is greater than the average across all treatment and replicate (n=2).

The maximal growth rate data of *E. coli* presented on Fig. 4a) was used to determine the value of *ψ* and Φ_*threshold*_ in equation (25) and simulate the whole osmolarity-related growth curve (Fig. 4b). By doing so, we can qualitatively assess the accuracy of predictions. Φ_*threshold*_ was set 0.1643 mol/L, the Φ value leading to the highest growth rate and *ψ* was set to 1020 to reproduce the negative slope of 3.8 *L* × *mol*^−1^ × *h*^−1^ found experimentally. On the hypo-osmotic side of the curve, both the experimental and computational results show that growth rapidly increases with osmolarity. Furthermore, in both cases, growth initially increases sharply and then modestly as osmolarity increases further. In terms of differences, the rates of improvement are dramatically higher in the simulations. However, this should be taken with a grain of salt, as Φ is difficult to calculate *in vivo*, requiring full determination of the metabolome. We can nonetheless see that the effect of osmolarity on growth predicted by ocFBA qualitatively matches the experimental measurements. On the hyper-osmotic side, given that the experimentally measured growth appears to decrease linearly, the pseudo reaction reproduce the effects of high osmolarity on growth.

For our second set of validations, the same growth assays were performed using four different strains from the Keio collection [31]: JW1693 (ΔydiA), a control strain containing a deletion which minimally affect growth, JW1840, a strain containing a deletion for 6-phosphogluconate dehydratase (EDD), the first upstream reaction involved in the ED pathway (ΔEDD), JW2011 (ΔGND), a strain containing a deletion for phosphogluconate dehydrogenase (GND), the first upstream reaction involved in the PP pathway, and JW0750 (ΔPGL), a strain containing a deletion for 6-phosphogluconolactonase (PGL), the first upstream reaction involved in both the PP and the ED pathways. The goal of this experiment was to evaluate the predictions presented in Fig. (3). According to the analysis, at low osmolarity, glycolysis and the citric acid cycle are not fully feasible. In turn, we hypothesize that cells would rely further on the PP and ED pathways at low osmolarities to produce biomass. If this is correct, the Keio mutants that cannot rely on the PP or the ED pathway should grow slower than the control at low osmolarities. JW1693 (ΔydiA) was chosen as the control instead of MG1655 because it contains the same added kanamycin resistance gene and was subjected to the same knock-out method as JW1840, JW2011 and JW0750. To improve comparability between each plate, the control strain was grown on the same plate as the test strains. The average growth rate of the test strain was divided by that of the control strain to normalize the data. The normalized growth results show that strains containing knock-outs limiting the metabolic flows into the PP pathways (ΔGND and ΔPGL) performed significantly worse (p *<* 0.05, two-sided, from Kruskal-Wallis test, see Table S2 for test statistics) than the control strain when at least 0.04 mol/L of NaCl was added to the media (theoretical osmolarity of 153 mOsm) (Fig. 4c). The ΔEDD strain did not appear to suffer from growth disturbance for any of the tested conditions. This result suggests that down regulating the ED pathway exclusively does not impair growth for any osmolarities. In fact, with 0.02 and 0.04 mol/L of NaCl, the ΔEDD strain outperformed the control. However, the result also suggest that disturbing both the ED and PP pathway leads to greater growth defect at lower osmolarities, as the ΔPGL strain grew slower than ΔGND when 0.04 and 0.06 mol/L of NaCl was added to the media. Across all the tested salt concentrations, the addition of 80 mmol/L of NaCl (theoretical osmolarity of 233 mOsm) led to lowest normalized growth rate in all strains.

According to the ocFBA’s predictions, ΔEDD, ΔGND and ΔPGL should grow slower than the wild-type (WT) at both low and high osmolarities and growth defect should be greater at low osmolarities (Fig. 4d). The predicted growth of the ΔPGL strain is worse than that of the ΔGND strain which itself is worse than that of the ΔEDD strain. The predicted growth of ΔEDD is around 99% of the WT’s growth rate from Φ =0.1459 onwards. The predictions also show that upon reaching a non-limiting values on Φ, the ΔGND and ΔPGL will grow at the same rate.

Two major differences can be found when comparing the ocFBA’s predictions to the experimental validation: experimentally, growth defects are much greater at higher osmolarities for ΔGND and ΔPGL, and the growth rate defects do not appear until the media’s theoretical osmolarity reaches 0.153 mOsm. These differences suggest two things. First, there exists a phenotype at very low osmolarities (73 to 113 mOsm) which is not captured by ocFBA, and *E. coli* relies further on the PP pathway in optimal conditions than what FBA predicts. This result is to be expected, as it was reported that the usage of the PP pathways tends to increase with the growth rate [32]. At low osmolarities, we observe that the Keio strains all have a significant growth defect resulting in normalized growth rates being close to 1 for the PGL, EDD and GND deletions. In contrast, the computational predictions do not reflect this result. Such a discrepancy could be because under these conditions, the PP and the ED pathways are infeasible. As a result, the strains might rely on other unidentified metabolic pathways causing them to have similar growth rates.

Outside of these differences, the predictions show consistency with the experimental results. The computational results correctly predicted that ΔPGL would be the slowest growing strain, that ΔEDD would be minimally affected by osmolarity, that ΔPGL would grow slower than ΔGND at low osmolarity and similarly at the highest osmolarities, and that the growth defect across all species would be greater at low osmolarities. Overall, this validation added some nuance to the predictions, as the experimental results show that the PP pathway remains useful even at higher rates of growth. However, they did confirm that in hypo-osmotic environments, not being able to rely on the ED and the PP pathway leads to significantly lower rates of growth, as shown by the Δ*PGL* strain.

For our third set of validations, to confirm that osmolarity affected the total pool of metabolites, a metabolome analysis was performed on 10 cultures grown in 500 mL bioreactors (full list and peak area measured for all metabolites available on GitHub, see Data availability). Each bioreactor was operated in duplicates. A total of 5 treatments were evaluated. *E. coli* MG1655 was grown in our low osmolarity media supplemented with either 0.001 mol/L, 0.025 mol/L, 0.050 mol/L, 0.1 mol/L or 0.4 mol/L (theoretical osmolarity of 75 mOsm, 123 mOsm, 173 mOsm, 273 mOsm, 873 mOsm), to capture growth in low, optimal and high environmental osmotic pressure.

Calculating the exact size of the metabolome can be difficult given that some metabolites, such as ATP and NADH can rapidly interact with other components in solution [33]. Thus, a qualitative approach was adopted to compare each metabolome. Treatments were ranked based on the number of times each treatment led to higher metabolite concentration. We called this measure the abundance index (see Methods for full description). A high abundance index implies that the treatment often leads to a greater metabolite concentration.

According to our analysis, the treatment with 0.1 mol/L of NaCl had a higher abundance index than any other treatment (Fig. 4e). This was followed by the 0.4 mol/L treatment, the 0.05 mol/L treatment, 0.025 mol/L treatments and finally the 0.001 mol/L treatment. Judging from the growth rate associated with each treatment (Fig. S3), the abundance index cannot be attributed solely to faster growth, as even though the high osmolarity media resulted in the lowest rates of growth it scored second in terms of abundance. These results could imply that the metabolite abundance tends to increase with the media’s osmolarity. It should be noted that high osmolarities tend to be associated with higher concentrations of specific osmolytes [3, 4], meaning that the index associated with the 0.4 mol/L might under-estimate the total metabolite abundance.

In our computational analysis, growth phenotypes were found to be associated with specific metabolites involved in thermodynamic bottlenecks. To this effect, metabolites whose range across treatments varied by at least a factor of 5 were plotted (Fig. 4f). Among the most notable results, we find that the concentration of intermediates associated with PGL, GND, and EDD (ribulose 5-phosphate and 6-phosphogluconate) was higher in low osmotic conditions (0.001 mol/L and 0.025 mol/L of NaCl), while the concentration of intermediates associated with the citric acid cycle (pyruvate and phosphoenolpyruvate) was more abundant at the osmolarity associated with the highest growth rate (0.1 mol/L of NaCl). These results corroborate our predictions concerning the increased reliance on these pathways at low osmolarity. Additionally, we found that hypertonic media led to higher concentrations of reduced glutathione and of a disaccharide, potentially maltose or trehalose. Glutathione is known to play a role in the maintenance of high turgor pressure in hyperosmotic environments [5, 34], and trehalose is a known osmoprotectant [35].

It was found that alanine was about 4 times more concentrated in our lowest osmolarity media and that acetyl-CoA, was about 4 times more concentrated in the media supplemented with 0.1 mol/L of NaCl (Table 2). According to the computational analysis, alanine is never involved in any thermodynamic bottlenecks. Thus, the link between alanine and osmolarity is unclear. However, alanine can be used as a carbon and nitrogen source and its utilization as a carbon source has been associated with harsher growth conditions in biofilms [36]. As for acetyl-CoA, the model had predicted that, as it is involved in thermodynamic bottlenecks at high growth rates, it should be more concentrated at higher Φ values (Fig. 2d). We thus find a good fit between the experimental data and the predictions for this metabolite.

**Table 2.** Intracellular concentration of metabolites in *E. coli* grown in a low osmolarity media supplemented with NaCl.

| NaCl<br>(mmol/L) | Metabolite concentration (mmol/L) |  |  |  |  |  |
| --- | --- | --- | --- | --- | --- | --- |
|  | Acetyl-CoA | Alanine | Fumarate | Glutamate | Glutamine | Valine |
| 1 | 0.130 ± 0.012 | 10.727 ± 3.504 | BDL<sup>-1</sup> | 82.766 ± 2.995 | 1.564 ± 0.335 | 2.505 ± 0.273 |
| 25 | 0.705 ± 0.116 | 2.724 ± 0.056 | 0.206 ± 0.046 | 56.579 ± 6.066 | BDL<sup>-1</sup> | 4.671 ± 1.959 |
| 50 | 0.461 ± 0.120 | 3.218 ± 1.041 | BDL<sup>-1</sup> | 88.881 ± 1.119 | 2.389 ± 0.068 | 1.578 ± 0.059 |
| 100 | 2.537 ± 1.460 | 2.919 ± 0.429 | 0.316 ± 0.101 | 82.554 ± 45.345 | 3.198 ± 2.058 | 8.126 ± 3.203 |
| 400 | 0.136 ± 0.012 | 1.596 ± 0.413 | 0.162 ± 0.010 | BDL<sup>-1</sup> | BDL<sup>-1</sup> | BDL<sup>-1</sup> |

The value given is the midpoint between the concentration in both reactors, while the uncertainty is the distance between the midpoint and the concentration in each replicate (n=2)
<sup>1</sup>BDL means below the detection limit. The tag was given if at least one replicate satisfied this condition

The concentration of acetyl-CoA, alanine, fumarate, glutamate, glutamine and valine were in the same order of magnitude (same power of 10) as the ones reported in Bennet et al., 2009. This consistency is despite these results being measured with another *E. coli* strain and despite growth in a different media. However, the concentrations of fructose 1-6 bisphosphate, 6-phosphogluconate, citrate, G3P, G6P and malate were found to be below the concentration of our lowest standard, indicating possible variation in the central carbon metabolism metabolites. It is thus possible that some metabolites, such as amino acids which are not involved in other metabolic functions, maintain relatively stable concentrations regardless of conditions.

Glutamate was not found to accumulate at higher osmolarities. Although glutamate is often associated to adaptation to high osmolarity, it tends to be accumulated in the early stage of adaptation to osmotic stress. Later on, glutamate and potassium ions are replaced by glutathione and trehalose [5, 37].

## Discussion

### Growing under osmotic stress

No matter the conditions, some level of metabolic activity is required for survival, if only to counteract the entropy that the degradation of complex structures produced by living organisms could generate. Osmotic pressure can pose significant challenges, as it forces cells to adopt a growth phenotype that utilizes metabolites sparingly.

Through our *in silico* analysis as well as our *in vivo* results, we have found that osmotic pressure can severely impair growth. We argue that, as osmotic pressure limits the total concentration of metabolites, the total pool of feasible reactions diminishes. In turn, the basal metabolite concentration required to make growth-related reactions thermodynamically feasible cannot be met. Osmotic limitations thus force cells to adopt a metabolome efficient phenotype, as opposed to the carbon-efficient phenotype.

As the environmental osmotic pressure increases, the metabolome changes. Metabolites, such as DHAP and phosphoenolpyruvate, that played little to no role become abundant. Behind these changes are thermodynamic bottlenecks restricting the cell’s ability to produce biomass efficiently. Being unable to meet the metabolic demand at low osmolarities, cells must rely on other enzyme to grow efficiently. However, as osmolarity increases, the metabolites associated with these bottlenecks can be accumulated, leading to changes in the metabolome. We have found that even reactions with negative standard Gibbs free energy values can become limiting when the metabolites involved in the reaction are required in high concentrations for other reactions to occur.

Together, these constraints were highlighted through our computational and experimental observations. We have found that as cells are forced into metabolically conservative pathways, they must rely on metabolite-efficient yet stoichiometrically inefficient pathways to generate cofactors. According to our computational predictions, at lower smolarities, cells are forced to rely on the Entner-Doudoroff (ED) and the pentose phosphate (PP) pathways. As osmolarity increases, the full citric acid cycle becomes available, and eventually glycolysis can proceed in the forward direction.

Our research question was aimed at understanding why the growth rate of *E. coli* and other bacteria were systematically lower at steady-state in low and high osmolarity environments. We hypothesized that, in hypo-osmotic conditions, the pool of usable metabolites is reduced, making several reactions thermodynamically infeasible.

Through our ocFBA, we were able to accurately replicate the major trends of our experimental results. Through our abundance index, we were also able to support the predictions that the total concentration of metabolite cannot be solely attributed to the growth rate and that the media osmolarity does indeed increase the size of the metabolome.

To our knowledge, there are very few studies done on the steady-state metabolism of bacteria following osmotic stress and even fewer on hypo-osmotic stress. This phase is described as a “?” in the 1999 review from Wood [38]. Upregulation and downregulation of different genes following osmotic downshift have long been reported in the literature, but only in the context of preventing cell lysis following a rapid osmotic upshift of downshift [7, 39]. These genes tend to be associated with transporters used to export, in the case of osmotic downshift and import, in the case of osmotic upshift, rather than on the metabolism itself. One of the few comparable studies was done with the red algae *Gracilaria changii*. In this study, the algae was grown in medias with different level of salinity to find genes that were down- or up-regulated by hypo- or hyper-osmotic stress. Changes in gene expression was measured through changes in the RNA. Their study showed that fructose-bisphosphate aldolase (FBA) was down-regulated at low osmolarity and pyruvate kinase (PYK) was upregulated at higher osmolarity [40]. Our results tend to corroborate such findings, as we have found that, at low osmolarity, *E. coli* would rely less on glycolysis to generate biomass and that FBA could not proceed in the forward direction.

On the metabolome side, almost all previous research has focused on hyperosmotic stress. On that side of the curve, our experimental results matches those reported, as we have found increased concentrations of reduced glutathione and disaccharides in those conditions. However, as our model relies strictly on thermodynamics, the abundance of these osmoprotectants could not be predicted by ocFBA. On the hypo-osmotic of the curve, the literature is much less detailed. In the few cases where the hypo-osmotic effect of the growth metabolism was evaluated, results point in the same direction as our predictions and our experimental results, showing lower concentrations of specific metabolites in hypo-osmotic environments [41, 42].

In this study we developed a novel modeling framework and used it to systematically analyze the impact of osmolarity on metabolism. The study led to a mechanistic understanding of the effect of osmolarity on metabolism and highlighted the importance of thermodynamic bottlenecks brought about by osmolarity constraints. We also validated these predictions through a comprehensive metabolomic and physiologic studies. As every living organism must adapt to osmotic stress, some of its conclusions also extend beyond *E. coli*. In addition, we expect that the ocFBA can be used to understand the impact of osmolarity during overproduction of biochemicals in metabolic engineering and synthetic biology applications. Improved mechanistic understanding of the effect of osmolarity and associated models will enable the design of robust strains for scaling up bioprocesses for commercial production.

## Materials and methods

### Metabolic steady-state constraints

Our formulation is built upon the main assumption behind FBA, which is that reaction fluxes are balanced during growth at steady-state. Thus, our first set of constraints is identical to FBA: the formulated problem contains a stoichiometric matrix **N** with the number of moles of every reactant and product involved in each reaction, a vector **r** representing the flux of each reaction and a set of upper (*β*_*i*_) and lower (*α*_*i*_) bounds constraining fluxes to biologically feasible rates. These sets of constraints can be written as

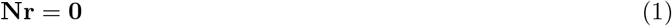

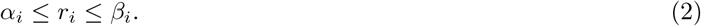

### Metabolic model

The most recent genome-scale *E. coli* stoichiometric model, *i* ML1515 [22], was used in our formulation. Every reversible reaction in *i* ML1515 was split into a forward and a reverse reaction to simplify the formulation. The modified model consisted of 3683 reactions and 1877 metabolites. Like *i* ML1515, the maximum glucose uptake rate was set to 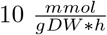 and all other carbon source uptake reactions were blocked.

### Thermodynamic constraints

At constant pH, the Gibbs free energy change of each reaction 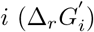 can be calculated according to the formula

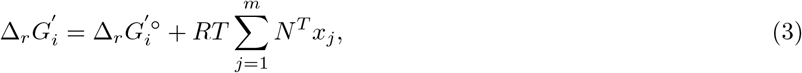

where 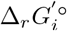 is the Gibbs free energy change of reaction *i* in standard conditions, *R* is the gas constant (kJ*· K*^−1^*· mol*^−1^), *T* is the temperature in Kelvin and *x*_*j*_ is the log of the concentration of metabolite *j* (*x*_*j*_ = *ln*[*C*_*j*_]) involved in reaction *i* [14].

To ensure that reactions with a non-zero flux in the solution are thermodynamically feasible, a binary variable *k*_*i*_ *∈ {*0, 1*}* was introduced and two subsequent constraints, inspired by OptMDFpathway [20] were formulated:

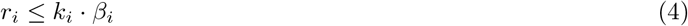

and

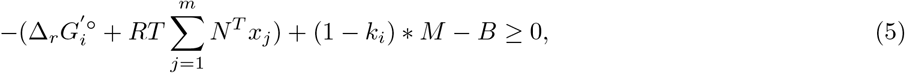

where

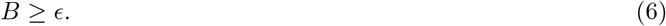

Here, M is a very large number, *B* is the minimal driving force of all active reactions and *ϵ* is the chosen minimal driving force value.

The minimal and maximal standard metabolite concentrations were bounded according to physiological data [43, 44]. The lower bound (*C*_*min*_) was fixed to 10^−6^ M and the upper bound (*C*_*max*_) was set to 0.1 M. This limitation can be formulated as the constraint

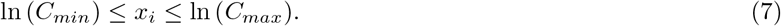

The concentration of water and protons H^+^ was set to 1M in the problem. The concentration of those two components is always set to 1M in eQuilibrator. For (H^+^), this is because its concentration is already accounted for, as the pH is used to determine 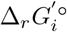 Similarly, for water the concentration is also already accounted for.

### Osmotic balance constraint

Osmosis affects every electrolyte system where there is a difference in the solute concentration between two environments separated by a membrane. For bacteria, a difference in ion concentration between the outside environment and the cytoplasm creates an osmotic pressure that drives water passively in or out of the cell.

Assuming water is the main solvent inside and outside the cell, the osmotic pressure can be estimated with the van’t Hoff formula:

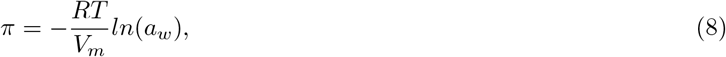

where *π* is the osmotic pressure in bar, *R* is the gas constant, *T* is the temperature in kelvin, *V*_*m*_ is the molar volume of water in mol m^-3^ and *a*_*w*_ is the water activity.

In environments with multiple ions, the water activity (*a*_*w*_) can be calculated by taking the product of each water activity (*a*_*w,j*_), using the Ross equation [45]:

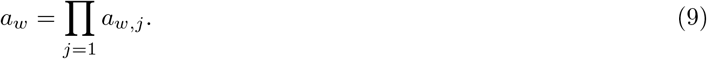

Ross’s equation assumes the absence of interaction between ions. The water activity of each metabolite *a*_*w,j*_ is defined as

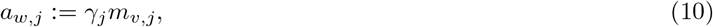

where *γ*_*j*_ is the free ion activity coefficient of metabolite *j* and *m*_*v,j*_ is the molar fraction of water in a solution containing the solute *j*. Note that *m*_*v,j*_ is equal to 1− *m*_*s,j*_, where *m*_*s,j*_ is the molar fraction of the solute *j* in solution.

For the free ion activity coefficient (*γ*_*j*_) calculation, we used Davies equation:

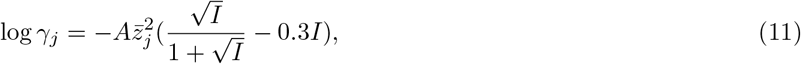

where *I* is the ionic strength, A is the dielectric constant of water, which is equal to 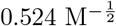 at 37^*°*^C [46], and 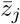 is the average charge of metabolite *j*. Davies equation is an empirical extension to the Debye-Hückel equation and is better suited for an environment in which ionic strength reaches 0.5M. Its wider applicability range makes it more accurate for the intracellular environment of *E. coli* where the ionic strength is estimated to fall in a range between 0.2 and 0.6M [47].

The ionic strength *I* can be reasonably assumed to be constant in the cell and is a function of the charge and concentrations of all metabolites [48]:

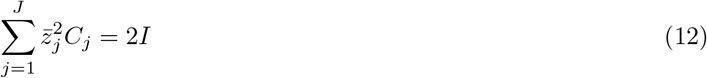

At this point our formulation for the osmotic pressure in the cytoplasm is

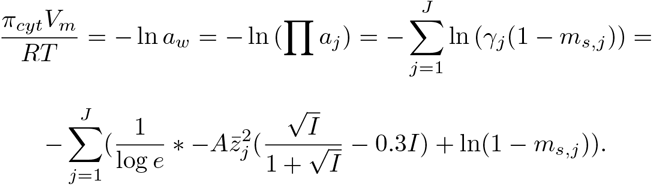

Note that, as *m*_*s,j*_ is very small, *ln*(1 − *m*_*s,j*_) *≈* −*m*_*s,j*_ and that the molar fraction of solute in a solution is defined as

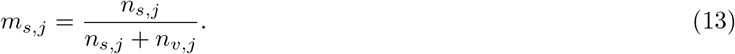

Since the number of moles of solute *j* in solution (*n*_*s,j*_) is much smaller than the number of moles of solvent (water) in solution (*n*_*v,j*_), we can approximate *m*_*s,j*_ as *n*_*s,j*_*/n*_*v,j*_.

Taking those into account, we find that the osmotic pressure is given by

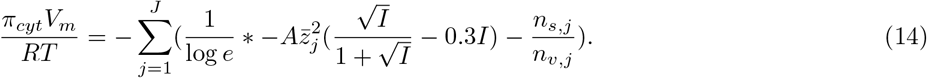

As

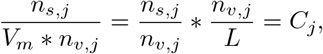

we can reformulate equation 14 as

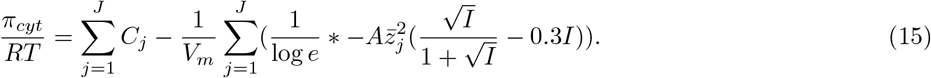

Using equation (15), we can determine the osmotic pressure of the extracellular environment by taking into account every solute *e*, allowing us to calculate the osmotic pressure of the cell in its environment instead of just pure water

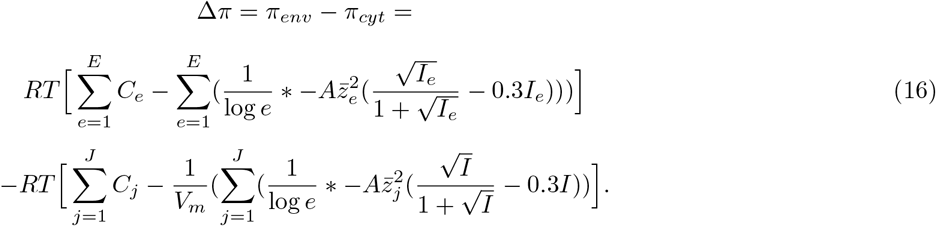

Here, to reduce the uncertainty associated with each constant, we introduced the osmotic concentration coefficient Φ, which units are mol/L and which incorporates the constants *π*_*env*_, *I, A* and 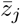, giving:

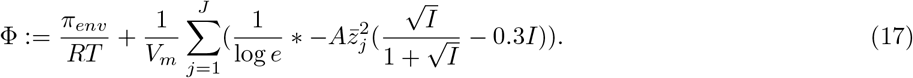

From this, we get

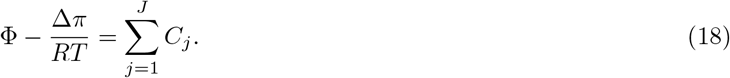

According to experimental data Δ*π* remains constant around -0.3 atm [21]. However, although cells have mechanisms that ensures pressure inside the cell remains negative, turgor pressure is not essential to growth [10]. Thus, we can introduce a new constant *θ* representing a lower limit for the osmotic pressure applied to the cell wall. As for the upper limit, to ensure pressure does not become positive, we set its value to 0 such that

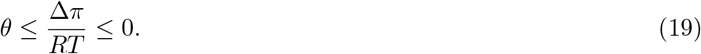

Given this information, we can pair both Δ*π* and Φ to represent the feasible intracellular concentration range, giving

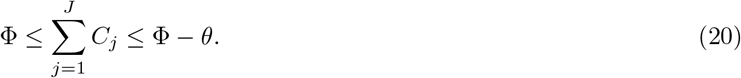

In all of our analysis, we allowed for the minimal value of Δ*π* of -0.3 atm and thus, *θ* was set to -0.01 mol/L (−0.3 atm /RT). This value can have a strong impact on the exact value of Φ associated with a specific growth rate but should have no impact on the dynamic of the relationship. This should be kept in mind when comparing Φ to experimental values.

This formulation departs from Akbari et al., 2021 work, which relies on Pitzer’s model to calculate water activity. Although to be completely accurate, Pitzer’s model should be used, it relies on many parameters which would either be impossible or extremely tedious to find in the context of biological growth. Thus, Akbari et al. made multiple assumptions which resulted in a set of constraints almost identical to ours, i.e. that the osmotic pressure depends on the sum of all metabolites and the ionic strength of the solution. The main difference in our formulation is the inclusion of the effective charge of each metabolite.

### Metabolite concentration approximation

At this point, we have an osmotic pressure constraint which involves the molar concentration (*C*_*j*_) of each metabolite and a thermodynamic constraint which depends on the natural logarithm (*x*_*j*_) of the concentration (*C*_*j*_) of those same metabolites: 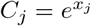. To relate those two together while preserving linearity, the variable 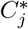, equal to 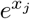, was linearly approximated through a piecewise linear function. We denote the metabolite concentration approximation as 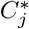 to prevent any confusion with the exact value. This approximation can be formulated as the constraint:

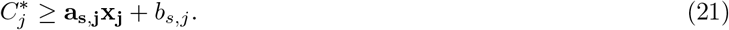

Here, a set *S* = *{*1, …, *n}* represents all the pieces of the function used to estimate *C*_*j*_ within the concentration range of 1*μ*M to 0.1 M, *a*_*s,j*_ is the slope of the function within section *s ∈ S* and *b*_*s,j*_ is the y axis shift of the function within this same interval. For each section *s* the slope *a*_*s*_ is given by

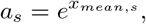

where 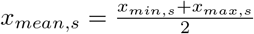. The variables *x*_*min,s*_ and *x*_*max,s*_ are the natural logarithm of the concentration interval in each piece *s*. The y-axis shift is given by

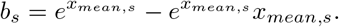

The minimal number of required pieces and the values of the intercepts were calculated by iteratively increasing the size of *s* until

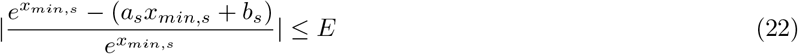

and

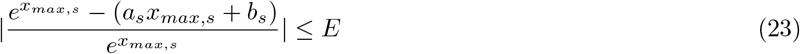

where *E* is the specified relative approximation error. For each interval, a new slope *a*_*s*_ and a new y-intercept *b*_*s*_ were calculated.

The value of *E* has the potential to exponentially increase the number of variables and constraints in the problem. The number of new variables can be calculated with

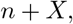

and the number of new constrains can be calculated with

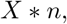

where *X* is the total number of metabolites whose concentration is estimated in the problem and *n* = | *S*|, determined by increasing the size of *S* until equation (22) and (23) are satisfied. In our formulation, *E* was set to 0.05, which resulted in *n* being 22. As *X* is 782, the linear approximation added 804 new variables and 17204 constraints to the problem. Despite *E* being 5%, when Φ remained in the range of 0.146 to 0.5 M, the maximal measured error was 3.67% (Fig. S1).

The linear approximation given by equation (21) becomes a *de facto* strict equality when *C*_*j*_ is constrained by Φ in the solution. In our formulation, over a certain range, the osmotic concentration coefficient (Φ) limits the total sum of all metabolite concentrations, keeping the approximation error below 5%. (Fig. 5). Note that, in these conditions we can get rid of the lower bound in equation (20), as *C*_*j*_ will always be as high as required.

**Fig 5.**
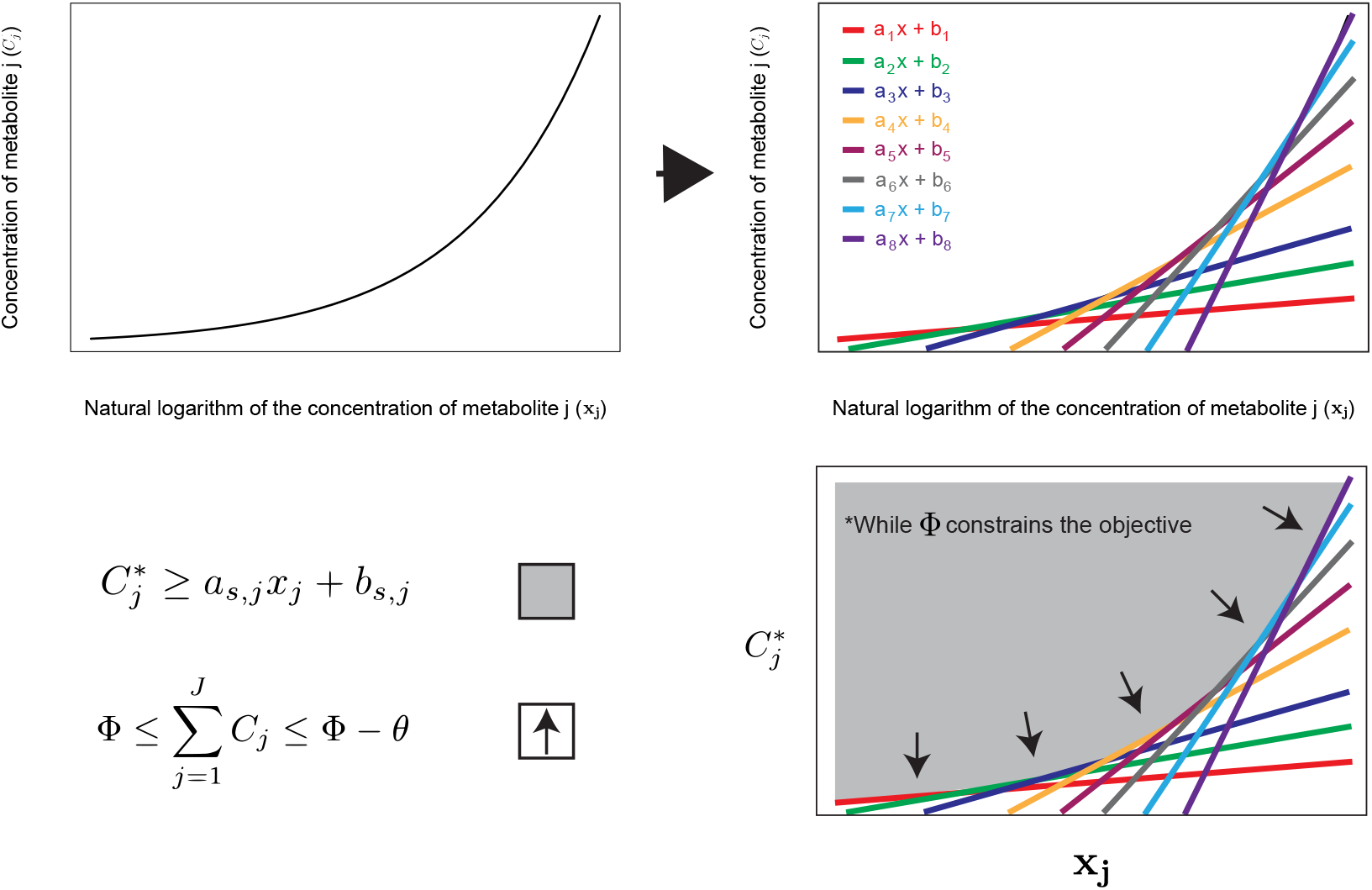
Linear approximation of the natural logarithm of the metabolite concentration through a piece-wise linear function. Through the formulation of *s* linear constraints, given the natural logarithm of the concentration, the concentration can be approximated for as long as 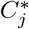 is minimized in the problem. Given equation (20), for as long as Φ limits the objective function of equation (26), 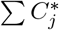 will be equal to Φ − *θ*. As the concentration of each metabolite matters and the space is limited, the problem will minimize 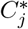, pushing its value in the solution along the edges of the polyhedron created by the *s* linear constraints, making the approximation accurate. The zone in grey represents the solution space.

### Hyper-osmotic constraint

The current state of the problem implies that past a certain Φ threshold, osmotic pressure no longer limits the pool of metabolome and thus, no longer affects growth. However, at high osmolarities, growth is known to decrease proportionally with the media’s osmolarity [42]. This phenomenon can be attributed to the cost associated with the active importation of ions, usually potassium, used to match the osmolarity of the media and to higher maintenance costs associated with the production of osmolytes [3–5]. To capture this increase in costs, we introduced a pseudo reactions, called Osmotic maintenance Demand (OmD) which consumes ATP proportionally to Φ. OmD duplicates the ATP maintenance (ATPM) reaction already in the model:

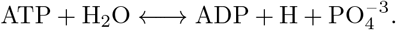

However, the rate of this reaction is constrained by *ψ*, the demand ratio (*mmol*× *L* × *mol*^−1^ × *g*− *DW*^−1^ × *h*^−1^) and Φ the environmental osmolarity proxy variable, such that

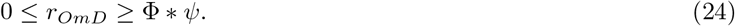

Note that it must remain positive to preserve its meaning.

To ensure that this demand only had to be met at high osmolarity, a threshold constant, Φ_*threshold*_, representing the values of Φ at which the media starts being hyper-osmotic, was added to our final formulation, giving

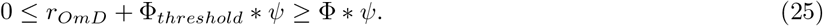

### Osmotically constrained flux balance analysis

Using equations (1), (2), (4), (5), (6), (7),(20), (21) and (25), we formulated a Osmotically Constrained Flux Balance Analysis (ocFBA) that allowed us to calculate the optimal *μ* for different osmotic concentration coefficient Φ:

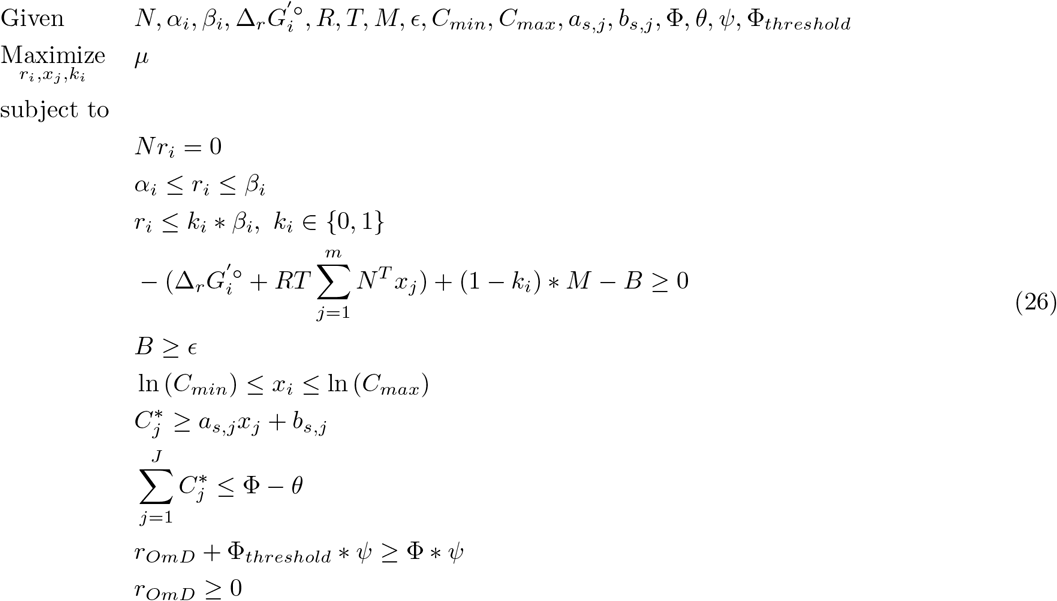

### Identification of thermodynamic bottleneck reactions

Thermodynamic bottleneck reactions can be defined as reactions that constrain the maximal value that *ϵ* could get. These bottlenecks were identified through successive optimizations involving modified versions of problem (26). Given *μ* and Φ, a first optimization finds the maximal value of *B* with

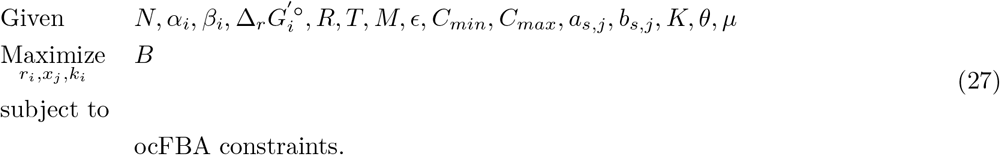

Then, reactions in the solution of problem (27) whose driving forces are equal to *B* are identified. Individually, the thermodynamic driving force of these reactions is set to 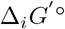. and problem (27) is solved again. If the new solution yields a higher value of *B*, the reaction is tagged as a bottleneck. This is done for each reaction in the set. This method allows the identification of single or multiple (distributed) bottleneck reactions.

### Concentration and Flux Variability Analysis (CVA and FVA)

The range of every feasible metabolite concentration was captured through a *Concentration Variability Analysis* (CVA) [14]. CVA can be formulated, given *ϵ*, Φ and *μ*_*max*_, by changing the objective function of equation (26) to

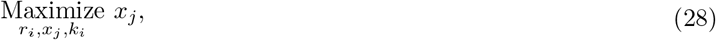

and then by

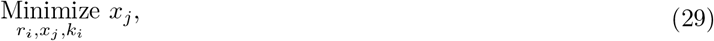

for every metabolite *j*.

Similarly, to evaluate the full range of feasible fluxes, we used Flux Variability Analysis (FVA) on all reactions *i* [49].

### Collection of thermodynamic data

We collected the change in Gibbs free energy of the system in standard conditions associated with each reaction Δ_*r*_*G*^*′°*^ for all single-compartment reactions using the eQuilibrator API [50]. This method requires a specified pH, pMg and ionic strength, these were set to 7.5, 2.5 and 250 M, respectively. These values are determined based on an adapted version of the group contribution method implemented in eQuilibrator by Noor *et al*. [51]. Multi-compartmental reactions were left thermodynamically unbounded. This was done because the calculation of their respective standard Gibbs free energies relies on membrane potential differences, a variable not considered in our analysis.

Upon solving equation (26) it was found that when *ϵ >* 0, growth is not possible. To ensure that all possible growth rates identified by FBA were thermodynamically feasible, bottlenecks making the maximal growth rate infeasible were identified using the method described above. These identified reactions were subsequently considered thermodynamically unbounded in our formulation. These reactions were sirohydrochlorin dehydrogenase (SHCHD2), 3-deoxy-manno-octulosonate cytidylyltransferase (KDOCT2), 2-C-methyl-D-erythritol 2,4-cyclodiphosphate synthase (MECDPS), aiaminohydroxyphosphoribosylaminopryrimidine deaminase (DHPPDA2), ATP phosphoribosyltransferase (ATPPRT), imidazole-glycerol-3-phosphate synthase (IG3PS), murein crosslinking transpeptidase (MCTP1App) and phosphoribosylaminoimidazole carboxylase (AIRC3). As a result, our analysis may underestimate the true concentration of the metabolites involved with these reactions.

### Uncertainty of thermodynamic data

To account for the uncertainty (*σ*_i_) associated with each Gibbs free energy values 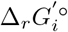 in equation (26) was set to 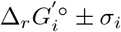. As a result, the growth curve is shifted towards the left or the right, but the overall shape remains unchanged (Fig. S3).

The bottleneck reactions remained similar as well, although some new ones appeared when considering the uncertainty on Δ*G*^*′°*^ (see full list in Supplementary Information). The reason for this change is tied to the value of Φ for which the bottleneck analysis is done. When uncertainty is considered, the same phenotype become feasible with smaller Φ values, resulting in a more constrained metabolome.

### Implementation

Python 3.8.13 was used for all calculations, expect for Figure 1 which was made with Matlab 2019b and Gurobi 10.0.2. COBRApy was used to handle metabolic model and perform flux balance analysis [52]. The Python library PuLP [53] was used to construct MILPs and IBM’s CPLEX 22.1 Python API was used to solve them.

### Experimental methods

We formulated a minimal low osmolarity media using the exchange reactions values of parsimonious FBA on *i* ML1515 [54]. The media is made of a sulfate solution containing 0.016 mM of FeSO_4_, 0.0014 mM of MnSO_4_, 0.0014 mM of CuSO_4_, a chloride salt solution containing 0.02mM of MgCl_2_, 0.0007 mM of ZnCl_2_, 0.01 mM of CaCl_2_, 0.0006 mM of NiCl_2_, 50 *μ*M of CoCl_2_, 7 *μ*M of Na_2_MoO_4_, a buffer and nitrogen solution containing 16.25 mM of K_2_HPO_4_, 3.75 mM of KH_2_PO_4_, 5.5 mM of NH_4_SO_4_ and a sugar solution containing 5.5 mM of glucose. Based on the formulation, the media should have a theoretical osmolarity (assuming full dissociation of ions) of 73 mOsm. The buffer concentration was experimentally determined by progressively increasing the concentration of K_2_HPO_4_ and KH_2_PO_4_ in the solution until the pH remained above 6.5 upon complete glucose consumption. For sterilization, the sugar solution was autoclaved, whereas the rest of the solutions were filter sterilized. To change the media’s osmolarity, a solution of autoclaved NaCl or KCl was added to the media until the desired concentration was reached.

All growth assays were performed using a Tecan’s Spark^®^ for the growth rate subjected to different salt concentrations and a Tecan’s Sunrise^®^ for the Keio strains experiment. The readings were done on Falcon^®^ 96-well Clear Flat Bottom TC-treated Culture Microplates. The first and last column and the last row of the plate were left empty to prevent variability between replicates. The first row of the plate contained the blanks associated with each treatment (media with different salt concentrations). Optical density at 600 nm (OD_600nm_) was taken every 15 minutes for each replicates, repeatedly. The plate was agitated (double orbital with an amplitude of 5 mm) between each measurement and the temperature was maintained at 37 ^*°*^C. Growth experiments lasted for 24 h and all cultures reached the stationary phase before the end of the assays. The number of technical replicates depended on test performed. Data collection was done using the manufacturer softwares (Magellan 7.5 or SPARKCONTROL).

For the growth rate analysis, a Python code was made to calculate the maximum growth rate automatically (available on GitHub) using a method inspired by Kurokawa and Ying, 2017 [55]. First, the OD measurements were converted into the natural logarithm of those values, then all the data was fragmented into groups of 20 subsequent points, representing 5h of measurement. The slope of these five hours fragments (max growth rate) was then calculated and the R^2^ of these linear fragments were calculated to see if the data fitted a linear regression. If the highest max growth rate of the fragments had an associated *R*^2^ *≥* 0.99, then this value was saved as the highest growth rate. If it was not, the algorithm would search for the second and third highest values, and if those had an *R*^2^ *≥* 0.99 they would be taken as the maximal growth rate value. If none matched the criterion, the number of subsequent points was reduced by 1 for a minimal range of subsequent values covering at least 2.5h of measurements.

The strain used depended on the experiment carried. *E. coli* MG1655 was used to test the effect of NaCl and KCl on growth. For the effect of specific knock-outs on growth, different strains from the Keio collection (Horizon Discovery) were used [31]. The strain JW1693 (ΔydiA) was used as a control in these experiments as it was not found to play a role in osmolarity and results in growth rates similar to the wild-type strain in a minimal media [56]. This control strain was grown alongside the test strain for each plate to normalize the data afterward. The other strains used were JW0750 (ΔybhE), JW1840 (ΔEDD) and JW2011 (ΔGND). JW0750 contains a deletion which removes 6-phosphogluconolactonase (PGL), an upstream reaction involved in both the ED pathway and the PP pathway. JW1840 contains a deletion which removes 6-phosphogluconate dehydratase (EDD) an upstream reaction involved in the ED pathway. JW2011 contains a deletion which removes Phosphogluconate dehydrogenase (GND), an upstream reaction involved in the PP pathway. Although flux through these pathways is technically possible, due to them being thermodynamically driven in the cell’s environment, it should be greatly limited by the deletions. Pre-cultures for these strains were done in our media supplemented with 25 *μ*g/mL of kanamycin.

The results of the knock-out experiment were normalized to minimize the variations associated to each plate using the measured growth rate of the control strain (JW1693)

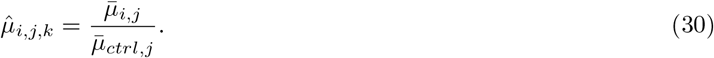

Here *μ*_*i,j,k*_ is the maximal growth rate of strain *i* ∈ {*ctrl*, Δ*GND*, Δ*PGL*, Δ*EDD*}, subject to a treatment *j* associate to replicate *k*. The variable 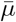 represents the average max growth rate. The uncertainty on the measured 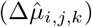 was calculated according to the empirical uncertainty calculation

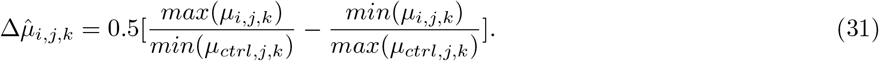

All statistical analyses were performed using R, version 4.2.2.

### Metabolome analysis

*E. coli* MG1655 cultures were grown in 500 mL Applikon MiniBio bioreactors in the low osmolarity media described above. The media was supplemented with 0.001, 0.025, 0.05, 0.1 and 0.4 mol/L of NaCl. The oxygen level of the culture was maintained above 30% through mixing and air sparging. The minimal stirring speed was set to 500 rpm to keep the media completely mixed. Cells were grown until OD_600nm_ 0.5. Subsequently, 20 mL of the fermentation broth was centrifuged for 3 minutes at 5000 rpm to collect the biomass. 1 mL of a chilled (−20 ^*°*^C) extraction solution containing methanol, acetonitrile and water (40:40:20) was added to the centrifuged biomass before incubation at -20 °C for 30 minutes. After incubation, the solution was centrifuged at 10,000 x g for 2 minutes. The supernatant was transferred to a tube and 0.3 mL of the extraction solution was added to the pellet and incubated at -20 ^*°*^C for 30 minutes before centrifuging again. After incubation, the solution was centrifuged, the supernatant was transfered to the same tube as earlier and the entire process was repeated once more using 0.2 mL of the extraction solution. Samples were then divided into two equal fractions before being concentrated in an Eppendorf Vacufuge plus speed vac and re-suspended in water. The volume of water added depended on the exact OD_600nm_ of the samples at the end of the fermentation. Each condition was tested in duplicate.

Untargeted mass spectrometry analysis was performed using an Ultimate 3000 UHPLC (Thermo Scientific) coupled to a mass spectrometer (MS) with an Orbitrap mass analyzer (Thermo Scientific Q Exactive) equipped with a HESI II probe. To get a wider range of metabolites, two columns were used: a Waters Atlantis Premier C18 Ax (1.7 *μ*m, 2.1 mm x 100 mm) and a Waters Acquity BEH HILIC (1.7 *μ*m, 3 mm x 150 mm) column (see further details in Supplementary Information). Compound identity was punitively assessed using the accurate mass of the parent ion and isotope patterns. Untargeted data processing was conducted in Compound Discoverer 3.2 (Thermo Scientific) using a 5ppm error allowance for compound identity against either the KEGG, HMDB, and BioCyc databases or mzCloud.

To infer the concentration of selected metabolites, three serial dilutions of media samples supplemented with 0.1 mol/L of NaCl and containing analytical standards (acetyl-CoA, aspartate, alanine, citrate, DHAP, fructose-6-bisphosphate, fumarate, G3P, G6P, glutamate, glutamine, malate, 6-phosphogluconate and valine) were processed through the same analysis method. The concentration of the analytical standards was determined based on the results from Bennet et al. 2009 (see Table S1 for concentration range). As four areas were assigned to the aspartic acid in our analysis and as all four had standard curves with a R^2^ greater than 0.9, it was not possible to differentiate the isomers enough to get a reliable concentration. It was thus excluded from our analysis.

The metabolite concentration per gram of dry weight was calculated according to

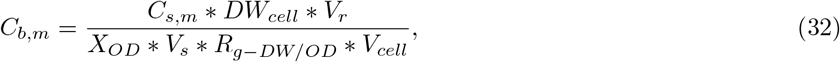

where *C*_*b,m*_ is the concentration of metabolite *m* in the cytoplasm of a cell in mmol/L-cytoplasm, *C*_*s,m*_ is the concentration of metabolite *m* in the sample analyzed in mmol/L-solvent, *DW*_*cell*_ is the dry weight mass of a single *E. coli* cell (2.8e-13g/cell from Neidhardt et al., 1996 [57], *V*_*r*_ is the volume of the concentrated re-suspension of the sample sent for MS analysis in L-solvent, *X*_*OD*_ is the OD of the bioreactors when cells were harvested, *V*_*s*_ is the volume of the sample (0.02 L), *R*_*g*−*DW/OD*_ is the ratio of cell dry weight per OD of *E. coli* (0.36 g-DW/(OD*L), from Ren et al., 2013 [58]) and *V*_*cell*_ is the volume of a single cell (6.7e-16 L-cytoplasm from Neidhardt et al., 1996 [57]).

### Abundance index

For all identified metabolites, each treatment was given a number from 1 to 5, according to the average height of its peak identified by mass spectrometry. A higher area means a higher concentration. However, these two variables are not necessarily linearly correlated, which is why an abundance index was designed for our analysis.

The index is calculated through subsequent steps:

1. find the peak area of each treatment and replicate for metabolite *i*;
2. for each treatment, find which replicate has the highest peak area;
3. assign a value from 1 to 5 to each treatments, according to the size of the respective area associated with the highest replicate (1 for highest area, 5 for lowest);
4. repeat the last step using the lowest replicate of each treatment;
5. save both list separately;
6. repeat step 1 to 5 for each metabolite and add the value to each lists.

At this point, 2 tables are generated, one for the high and one for the low replicate values. Using each table, the abundance index was calculated according to

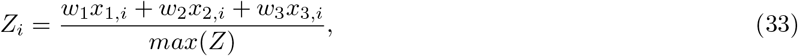

where *Z*_*i*_ is the abundance index associated with treatment *i, w*_1_, *w*_2_ and *w*_3_ are the weight associated with a rank of 1, 2 or 3, and *x*_1,*i*_, *x*_2,*i*_ and *x*_3,*i*_ are the sum of metabolites for treatment *i* being assigned the value of 1, 2 or 3, respectively. For weight *w*_1_, *w*_2_ and *w*_3_ were given the value 5, 3 and 1, respectively. Therefore, the maximal index value that could be given to a treatment (*max*(*Z*)) is equal to the total number of metabolites found by our putative analysis (414 with the HILIC column and 384 for the C18 column) multiplied by *w*_1_ or 5. The abundance index was calculated for table, giving a range with the lowest and highest possible index for each treatments.

## Data availability

All computational and experimental results used to make our figures are available on the LMSE GitHub page (https://github.com/LMSE/ocFBA).

## Code availability

All the codes used for our analysis are available on the LMSE GitHub page (https://github.com/LMSE/ocFBA).

## Supporting information

**Supplementary Information** Supplementary methods and results associated with the charge balance constraint, the metabolome analysis and the experimental validation experiment.

**Fig. S1 Accuracy of the concentration estimation**. Comparison between the approximated concentration values of 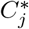 and the real concentration 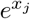 for different value of Φ. **a)** Comparison when Φ = 0.146*mol/L*. **b)** Comparison when Φ = 0.165*mol/L*. **c)** Comparison when Φ = 0.5*mol/L*. **d)** Comparison when Φ = 2*mol/L*

**Fig. S2 Effect of the uncertainty on the standard Gibb’s free energy value on the predicted growth rate for different** Φ **values**. The growth rate values were calculated for 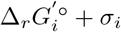 (red triangle), 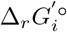 (black circle) and 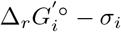 (green diamond).

**Fig. S3 OD of *E. coli* MG1655 grown in media with different osmolarities**. Cells were grown in bioreactors in our low osmolarity media until their population reached an OD of 0.5. The osmolarity of the media was adjusted by adding 0.001, 0.025, 0.05, 0.1 and 0.4 mol/L of NaCl, respectively. The reported theoretical osmolarity assumes full dissociation of all ions. Triangles and diamonds are used to differentiate between the two replicates.

**Table S1** Metabolite concentration in each of the standards used in the metabolome analysis

**Table S2** Test statistics for the Kruskal-Wallis test on the normalized growth rate of different strains

## Acknowledgements

We would like to acknowledge Robert Flick (Biozone, University of Toronto) for his help with the metabolomic sample preparation protocol and its subsequent analysis through mass spectrometry. Authors acknowledge funding from the Canada Research Chairs program and NSERC Discovery grant to R.M.

