## Supplementary Information for "Understanding systems level metabolic adaptation resulting from osmotic stress"

### Supplementary Methods

#### Metabolome analysis

The analysis of the metabolome on the fermentation samples was performed using an Ultimate 3000 UHPLC (Thermo Scientific) equipped with either a Waters Atlantis Premier C18 Ax (1.7 $\mu$m, 2.1 mm x 100 mm) column or a Waters Acquity BEH HILIC (1.7 $\mu$m, 3 mm x 150 mm), coupled to a mass spectrometer (MS) with an Orbitrap mass analyzer (Thermo Scientific Q Exactive) equipped with a HESI II probe. Mass spectrometry was performed using electrospray ionization under both positive and negative modes with fast polarity switching. MS1 spectra were acquired over an m/z range from 70 to 1050 with the mass resolution set to 70k, AGC Target of 3E6, max injection time 100 ms, spray voltage 3.5 kV, capillary temperature 320°C, sheath gas 40, aux gas 20, spare gas 5, and s-lens RF level 55. MS2 spectra were acquired using a Top10 method with a mass resolution of 17.5k, AGC Target of 1e5, max injection time of 50 ms, isolation window of 0.4 m/z and HCD collision energy of 30. For both columns, the oven temperature was set to 40°C.

For the C18 column, the eluents were composed of 0.1% formic acid in $H_2O$ (eluent A) and 0.1% formic acid in acetonitrile (eluent B). Gradient elution with a flow rate of 0.3 mL/min was performed at 5% eluent B for the first minute and increased linearly to 98% B from 1 to 7 min, held at 98% B from 7 to 10 min, then decreased linearly to 5% B from 10 to 10.5 min and held at 5% B until 15 min.

For the HILIQ column, the eluents were composed of 20mM Ammonium Acetate and 20mM Ammonium Hydroxide in $H_2O$ (eluent A) and acetonitrile (eluent B). Gradient elution with a flow rate of 0.4 mL/min was performed at 50% eluent B for the first minute and increased linearly to 100% A from 1 to 6 min, held at 100% A from 6 to 11.5 min, then decreased linearly to 50% B from 11.5 to 12 min and held at 50% B until 15 min.

Untargeted data processing was conducted using Compound Discoverer 3.2 (Thermo Scientific) using a 5ppm error allowance for compound identity against either the KEGG, HMDB, and BioCyc databases or mzCloud.

Some molecules with different formulas were assigned the same name by our putative analysis (potentially different protonation states). Each of them was treated as a different independent metabolite in our abundance index analysis. However, if that molecule was in our set of analytical standards (Table S1), only the formula for which the standard curve had a $R^2$ above 0.99 was kept to generate the heat map and calculate the concentration.

Table S1: Metabolite concentration in each of the standards used in the metabolome analysis

| | Concentration (mmol/L) | | |
|---|---|---|---|
| Metabolite | Standard 1 | Standard 2 | Standard 3 |
| 6-phosphogluconate | 5 | 0.5 | 0.05 |
| Acetyl-CoA | 1 | 0.1 | 0.01 |
| Alanine | 10 | 1 | 0.1 |
| Aspartate | 10 | 1 | 0.1 |
| Citrate | 5 | 0.5 | 0.05 |
| DHAP | 1 | 0.1 | 0.01 |
| F6bp | 25 | 2.5 | 0.25 |
| Fumarate | 1 | 0.1 | 0.01 |
| G3P | 1 | 0.1 | 0.01 |
| G6P | 10 | 1 | 0.1 |
| Glutamate | 200 | 20 | 2 |
| Glutamine | 10 | 1 | 0.1 |
| Malate | 5 | 0.5 | 0.05 |
| Valine | 10 | 1 | 0.1 |

### Supplementary Results

#### Accuracy of estimations

According to our approximation method, the piece-wise linear function should hold as long as $\Phi$ remains limiting. To ensure that the value of the approximation remained accurate for all of our analysis, we evaluated how the approximated concentration $C_j^*$ changed with the actual concentration $e^{x_j}$ for different $\Phi$ (Fig. S1). The average error associated to each metabolite is 0.07%, 0.14 %, 0.11% and 105,854% when $\Phi = 0.146$, $\Phi = 0.165$, $\Phi = 0.5$ and $\Phi = 2$ mol/L, repectively. The maximal error associated to each metabolite is 3.67%, 3.67%, 3.66% and 82672240 % when $\Phi = 0.146$, $\Phi = 0.165$, $\Phi = 0.5$ and $\Phi = 2$ mol/L, repectively. Interestingly, for $\Phi = 2$ mol/L, all the error is caused by a single metabolite, 2,3-Dihydroxy-3-methylbutanoate. Removing it makes brings the average and maximal error to 0.08% and 3.34 %. We thus find that the range of $\Phi$ chosen for this study (0.146M to 0.165) results in accurate approximations.

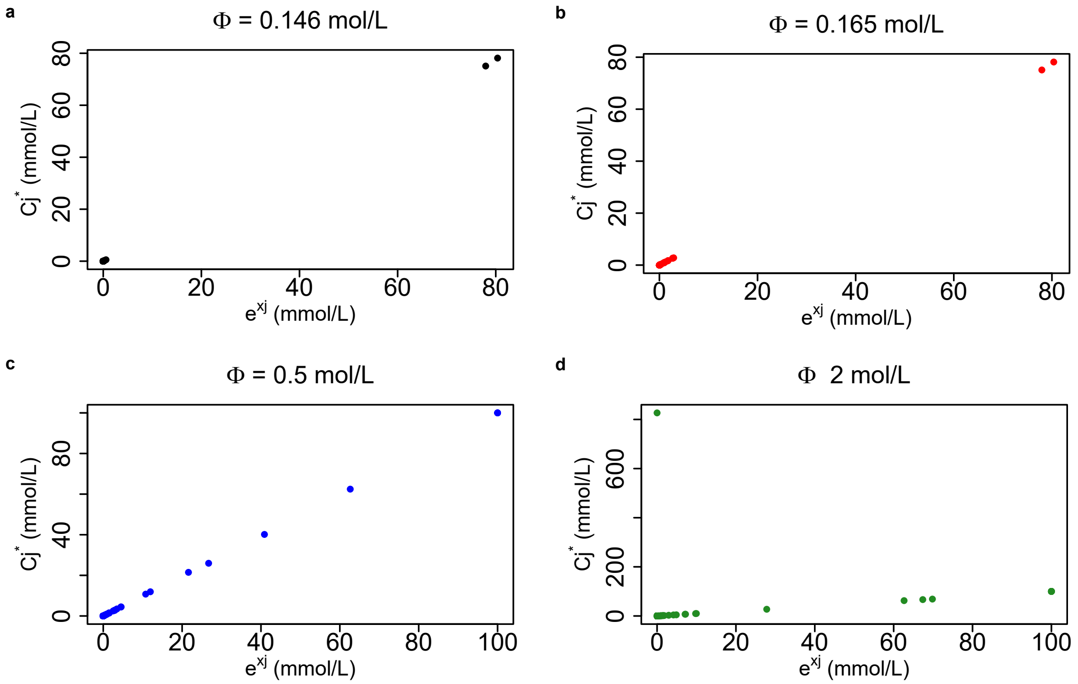


Figure S1: Accuracy of the concentration estimation. Comparison between the approximated concentration values of $C_j^*$ and the real concentration $e^{x_j}$ for different value of $\Phi$. **a)** Comparison when $\Phi = 0.146 mol/L$. **b)** Comparison when $\Phi = 0.165 mol/L$. **c)** Comparison when $\Phi = 0.5 mol/L$. **d)** Comparison when $\Phi = 2 mol/L$

#### Uncertainty of thermodynamic data

The effect of the uncertainty on the standard condition Gibb's free energy values ($\Delta_r G_i^{'\circ}$) on the relation between the growth rate and $\Phi$ was evaluated by solving the OCFBA problem while setting the standard condition Gibb's free energy value to $\Delta_r G_i^{'\circ} \pm \sigma_i$ (Fig. S2). According to the results, the shape of the curve remains identical regardless of the uncertainty. When the uncertainty was added to $\Delta_r G_i^{'\circ}$, the minimal value of $\Phi$ required to have some growth diminished and when it was subtracted, it increased. Although counter-intuitive, this is due to reverse direction reactions. By increasing the $\Delta_r G_i^{'\circ}$ of a reaction in the forward direction, we lower its value in the reverse direction.

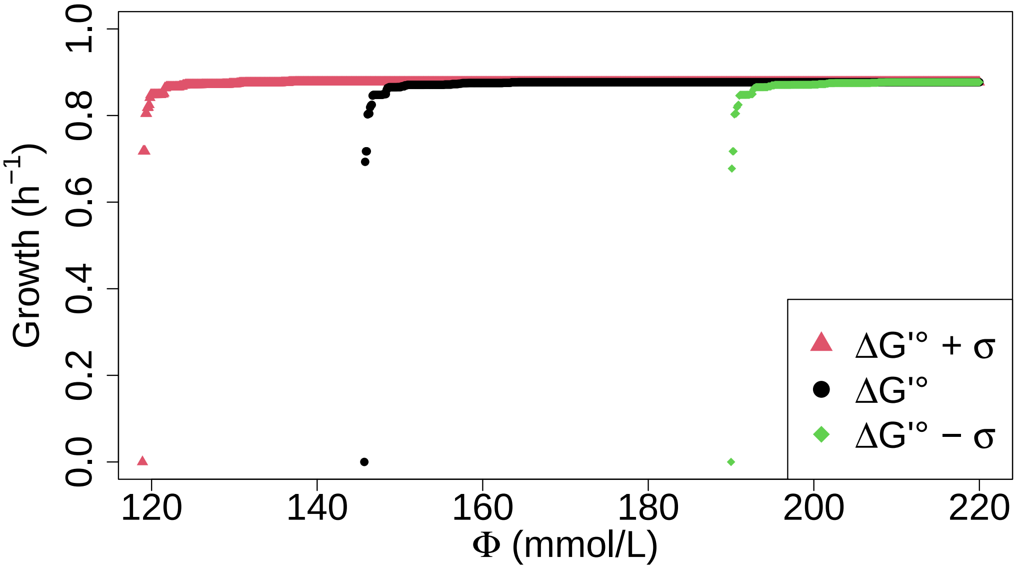


Figure S2: Effect of the uncertainty on the standard Gibb's free energy value on the predicted growth rate for different $\Phi$ values. The growth rate values were calculated for $\Delta_r G_i^{'\circ} + \sigma_i$ (red triangle), $\Delta_r G_i^{'\circ}$ (black circle) and $\Delta_r G_i^{'\circ} - \sigma_i$ (green diamond).

Some additional bottlenecks were found when considering the uncertainty on the standard condition Gibb's free energy values ($\Delta_r G_i^{'\circ}$). Without exception, all of those additional bottlenecks were found when $\Delta_r G_i^{'\circ}$ was set to $\Delta_r G_i^{'\circ} + \sigma_i$. For $\mu = 0.7174h^{-1}$, 2-isopropylmalate hydratase (IPPMIb, 0.3524 kJ/mol) and glutamate dehydrogenase (GLUDy, -33.0591 kJ/mol) in the reverse direction were found alongside the other bottlenecks in Table 1. For $\mu = 0.8045h^{-1}$, 3-isopropylmalate hydratase in the reverse direction (IPPMIa, 3.7575 kJ/mol), GLUDy in the reverse direction, phosphoglycerate mutase in the reverse direction (PGM, 4.1266 kJ/mol) and phosphoglycerate kinase in the reverse direction (PGK, -19.2273 kJ/mol) are now also bottlenecks. For $\mu = 0.8478h^{-1}$ it was fumarase in the reverse direction (FUM, -3.4279 kJ/mol), GLUDy in the reverse direction and AICART in the forward direction who were added. For $\mu = 0.8751h^{-1}$, aspartate-semialdehyde dehydrogenase in the reverse direction (ASAD, 24.316 kJ/mol), FBA3 in the forward direction, FUM in the forward direction, GLUDy in the reverse direction, PGI in the forward direction, PGK in the reverse direction, TALA in the reverse direction, AICART in the forward direction, phosphopentomutase in the reverse direction (PPM, 13.7023 kJ/mol) and ribose-5-phosphate isomerase in the reverse direction (RPI, -4.3534 kJ/mol) were added to the bottleneck list. For $\mu = 0.8770h^{-1}$, acetylglutamate kinase in the forward direction (ACGK, 25.7262 kJ/mol), ASAD in the reverse direction, ASPK in the forward direction, 3-hydroxyacyl-CoA dehydratase in the forward direction (ECOAH2, 0.1291 kJ/mol) and MTHFD in the forward direction we now bottlenecks.

#### Experimental validation

According to the non-parametric Kruskal-Wallis test on the growth rate of the control and test strain grown on the same plate, many of the treatment condition led to significantly different growth rate (see Table S2). The ΔEDD strain remains mostly similar to the control, whereas the ΔGND and the ΔPGL strains differ from the control when the NaCl is at least 60 mmol/L.

Table S2: Test statistics for the Kruskal-Wallis between different test strain and the control

| | Strain | | | | | |
|---|---|---|---|---|---|---|
| | ΔEDD | | ΔGND | | ΔPGL | |
| NaCl (mmol/L) | p-values | Chi-squared | p-values | Chi-squared | p-values | Chi-squared |
| 0 | 0.0495 | 3.8571 | 0.2752 | 1.1905 | 0.2752 | 1.1905 |
| 20 | 0.0495 | 3.8571 | 0.1266 | 2.3333 | 0.1266 | 2.333 |
| 40 | 0.0495 | 3.8571 | 0.0495 | 3.8571 | 0.0495 | 3.8571 |
| 60 | 0.2752 | 1.1905 | 0.0495 | 3.8571 | 0.0495 | 3.8571 |
| 80 | 0.8273 | 0.0476 | 0.0495 | 3.8571 | 0.0495 | 3.8571 |
| 100 | 0.8273 | 0.0476 | 0.0495 | 3.8571 | 0.0495 | 3.8571 |
| 120 | 0.8273 | 0.0476 | 0.0495 | 3.8571 | 0.0495 | 3.8571 |
| 140 | 0.2752 | 1.1905 | 0.0495 | 3.8571 | 0.0495 | 3.8571 |
| 160 | 0.5127 | 0.4286 | 0.0495 | 3.8571 | 0.0495 | 3.8571 |
| 180 | 0.0495 | 3.8571 | 0.0495 | 3.8571 | 0.0495 | 3.8571 |

To analyze the composition of the metabolome of *E. coli* MG1655 when grown in different osmotic conditions. Cells were grown in Applikon bioreactors in batch mode using our low osmolarity media (see Materials and Methods section).

The maximal growth rate of each bioreactor was calculated by finding the slope of the log value of $OD_{600nm}$ with respect to time for the time interval between the first and last hour of culture. The first hour of data was excluded to prevent misleading information associated with the lag phase of culture.

In accordance with the microplate experiments, the maximum growth rate was found to be highest in the media supplemented with 100 mmol/L of NaCl. ($0.61\ h^{-1} \pm 0.006$). This was followed by the media containing 50 mmol/L ($0.59\ h^{-1} \pm 0.002$), 25 mmol/L ($0.51\ h^{-1} \pm 0.026$), 1 mmol/L ( $0.43\ h^{-1} \pm 0.006$) and 400 mmol/L of salt ($0.30\ h^{-1} \pm 0.06$) (Fig. S3). One of the replicates associated with the 123 mOsm media reached OD 0.5 faster than all other treatments. However, this appeared to be associated with a particularly high rate of growth between 1 and 1.5h, which was not sustained throughout the culture.

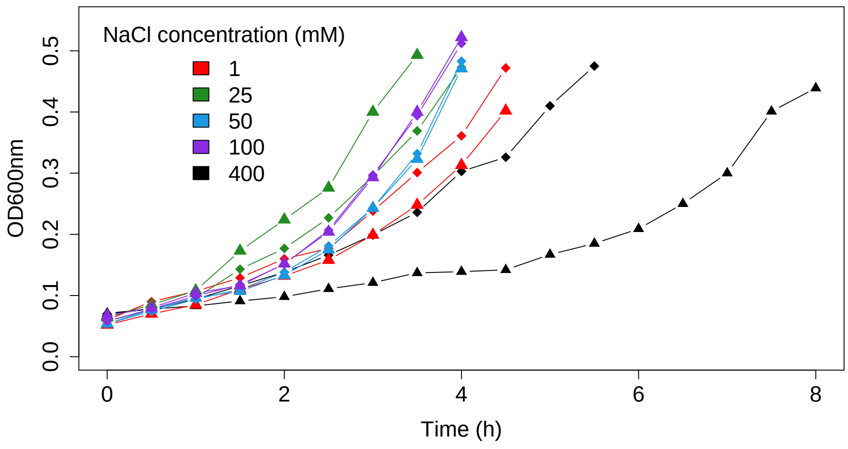


Figure S3: OD of *E. coli* MG1655 measured in bioreactors in media with different osmolarities. Cells were grown in bioreactors in our low osmolarity media until their population reached an OD of 5. The osmolarity of the media was adjusted by adding 1, 25, 50, 100 and 400 mmol/L of NaCl, respectively. The reported theoretical osmolarity assumes full dissociation of all ions. Triangles and diamonds are used to differentiate between the two replicates.

The abundance of intracellular metabolites was found to vary with each treatment (Table ??). The concentration of metabolite was found to be mostly higher in the culture growing in the media with 100 mmol/L of NaCl and consistently lower in the media with 400 mmol/L of NaCl. One notable exception is alanine which was about 4 times higher in the media with the lowest concentration of salt. The concentration of fructose 1-6 bisphosphate, 6-phosphogluconate, citrate, G3P, G6P and malate was found to be below the concentration of our lowest standard (0.25 mmol/L, 0.05 mmol/L, 0.05 mmol/L, 0.01 mmol/L, 0.1 mmol/L and 0.05 mmol/L, respectively). As our standards were prepared using data from Bennet et al., 2009 [1], values below the standard curve (BDL) means that the concentration was about 1.5 log smaller than the concentration reported in the paper.
